# A-LAVA: Detecting impact of germline variants on metabolic pathways in cancer genomes

**DOI:** 10.1101/2025.08.30.673276

**Authors:** Mansoureh Jalilkhany, Kiana Gallagher, Isabella Wehner, Phineas T. Hamilton, Sarah McPherson, Sarah McPhedran, Farouk Nathoo, Julian Lum, Ibrahim Numanagić

**Affiliations:** Department of Computer Science, University of Victoria, Victoria, B.C., Canada; Trev and Joyce Deeley Research, Centre, BC Cancer Agency, Victoria, B.C., Canada; Department of Statistics, University of Victoria, Victoria, B.C., Canada; Trev and Joyce Deeley Research, Centre, BC Cancer Agency, Department of Biochemistry and Microbiology, University of Victoria, Victoria, B.C., Canada

**Keywords:** metabolomics, gwas, gene set analysis, overlap correction

## Abstract

The metabolic landscape of cancer has been widely studied, especially in the context of somatic mutations. However, the impact of inherited germline variants upon the metabolic genes interaction still remains unexplored. In this work, we present a computational pipeline named A-LAVA for the detection and analysis of germline variants that affect metabolic pathways in cancer. Our pipeline enables analysis at three different levels: SNP, gene, and pathway-based analysis. The first steps consist of detecting statistically significant SNPs through standardized GWAS pipelines and, subsequently, genes associated with metabolic traits through gene-level analysis. Then, A-LAVA performs gene set analysis (GSA) to further explore the effect of detected associations on metabolic pathways. This analysis is done through a statistical model that newly corrects for the confounding effects arising from overlapping gene sets, in addition to other corrections performed by the current best practices. Our analysis conducted on TCGA data shows that SNP and gene-level results identified key associations and that A-LAVA’s GSA approach improved the overall accuracy both on synthetic and real data by correctly correcting for overlapping genes, refining significance thresholds, and reducing false positives, thus leading to more reliable metabolic pathway rankings and a more robust framework for gene set analysis.

**CCS CONCEPTS:**

- **Applied computing** →**Bioinformatics**; **Metabolomics / metabonomics**.

## 1 INTRODUCTION

Recent developments in cancer research enable us to detect, prevent or treat cancer diseases by uncovering genetic abnormalities that drive cancer development or resistance to treatment [4, 40, 46, 54, 57]. A key focus of these developments is the identification and classification of cancer-related somatic mutations that are directly linked to cancer diagnosis and recurrence and are, as such, directly targeted by various treatments. In addition, some genetic mutations can significantly alter treatment choices even when treatments do not directly target such mutations. For example, a mutation in the *TP53* gene means that cancer probably will not respond to chemoimmunotherapy in chronic lymphocytic leukemia [42]. Finally, the patient-specific cancer genome data enables us to advance precision medicine initiatives [19] by offering patients a more accurate diagnosis and, consequently, a more personalized treatment plan [42].

One important development is the study of cancer metabolism that reveals how altered metabolic processes support tumour growth, thus presenting new therapeutic targets [3, 12, 15, 22, 38, 47, 58]. The key assumption here is that, compared to healthy cells, cancer cells exhibit altered metabolic processes that contribute to developing and maintaining malignant characteristics [12]. Many studies focused on how changes in cell metabolism promote cancer cell survival and growth in various cancer types [3, 22, 38, 58]. Other studies focus on finding metabolic genes and processes that exhibit consistent changes in cancer cells across tumour types [47]. From a therapeutic standpoint, focusing on the metabolic variations between tumour and normal cells—i.e., *somatic mutations*— shows promise as a novel anticancer therapy [22, 37] (one such example is the study of the metabolic aspects of response to the immune checkpoint blockade therapy [15]). However, while the effect of somatic mutations on the metabolic landscape of cancer has been widely studied, the impact of the underlying germline (inherited) variants on the same landscape remains mostly unexplored [20]. Although some studies targeted the effect of germline variants in cancer [18, 34], they mainly investigated these effects on other phenotypes such as immune-related traits [48, 50]. As a result, germline variants and their impact on the interaction of metabolic genes as networks remain largely unexplored [18, 20, 34, 48, 50].

In order to address this gap, here we present a framework named A-LAVA for the discovery of the germline genetic variants and the associated genes and pathways that are correlated with metabolic-related traits. We began by utilizing genome-wide association study (GWAS)—a widely used methodology for finding disease or trait-related mutation sites [63]—on the large Cancer Genome Atlas (TCGA) RNA-seq dataset [61] to find the relevant germline variants that are correlated with metabolic gene set enrichment scores and, ultimately, various metabolic pathways.

Following the identification of statistically significant single nucleotide polymorphisms (SNPs) and genes, gene set analysis (GSA) methods were applied to examine the interaction of these associations. GSA integrates association signals across multiple genes within predefined gene sets, thus facilitating the interpretation of molecular mechanisms [59]. Nowadays, the commonly used tool for GSA in the GWAS context is MAGMA, which identifies significant gene sets associated with given traits while accounting for biases such as gene length and density [9, 49]. Other popular GSA analysis tools, such as DAVID [14, 51], GSEA [53], INRICH [30], GOATOOLS [28], or GOAT [29] are either not adjusted for the GWAS context (unlike MAGMA) or underperform in comparison to MAGMA [9].

One downside of MAGMA is that it does not adjust for overlapping gene sets—a common occurrence in metabolic datasets—and thus can lead to confounded associations [11]. To address this limitation, we present a new GSA model which refines MAGMA’s approach by correcting for shared genes across predefined gene sets. Our model reduces biases and improves the reliability of gene set rankings. As a result, A-LAVA’s outputs significantly altered *p* and *β* values that yield differing significance thresholds and rankings of associated gene sets as compared to MAGMA [11].

To evaluate A-LAVA, we first developed a controlled set of GSA simulations to show that it performs equally well or even better than MAGMA in the presence of overlapping gene sets. Then, we utilized KEGG (Kyoto Encyclopedia of Genes and Genomes)-defined gene sets [27] to calculate enrichment scores and conducted a GWAS study examining germline variants linked to 65 KEGG-based metabolic traits across 27 TCGA cancer cohorts representing ethnically diverse populations to identify key genes and pathways associated with metabolic traits in order to gain better insight into genetic interactions within metabolic networks. We detected 71 significant SNPs and 20 independent loci across 22 autosomal (non-sex) chromosomes for ten metabolic traits in the European ancestry. The most significant SNPs detected belonged to four metabolic traits of glutathione metabolism, drug metabolism (cytochrome P450), xenobiotics metabolism (cytochrome P450), and alpha-linolenic acid metabolism. For the pathway-level analysis, the top gene sets were identified for all metabolic traits using A-LAVA. We found a total of 201 significant pathways associated with 65 metabolic traits.

Overall, the presented results support our hypothesis that correcting for overlapping confounding factors results in more accurate *p* and *β*-values, and thus, in turn, improves final rankings of top gene sets. Thus, we hope that A-LAVA’s improved GSA modelling and enhanced detection toolkit will enable researchers to effectively utilize metabolomics data to further advance understanding of cancer prevention and treatment.

## 2 METHODS

A-LAVA is a framework for detecting germline variants that impact metabolic pathways and consists of the following steps (see Figure 1 for a graphical overview). In the first step, A-LAVA performs a genome-wide association study (GWAS) to identify germline SNPs that potentially influence human metabolic traits. Here, we began with germline variant data from TCGA [61] that was subsequently imputed with the missing SNPs to enhance genomic coverage [48]. To mitigate confounding genetic variations, samples were clustered by ancestry and went through rigorous quality control filters, such as minor allele frequency (MAF) thresholds and heterozygosity checks. Only unrelated individuals were retained for later analysis to ensure statistical robustness. Metabolic traits were inferred by processing TCGA RNA-Seq data and incorporating pathway enrichment scores to refine association signals [36].

**Figure 1:**
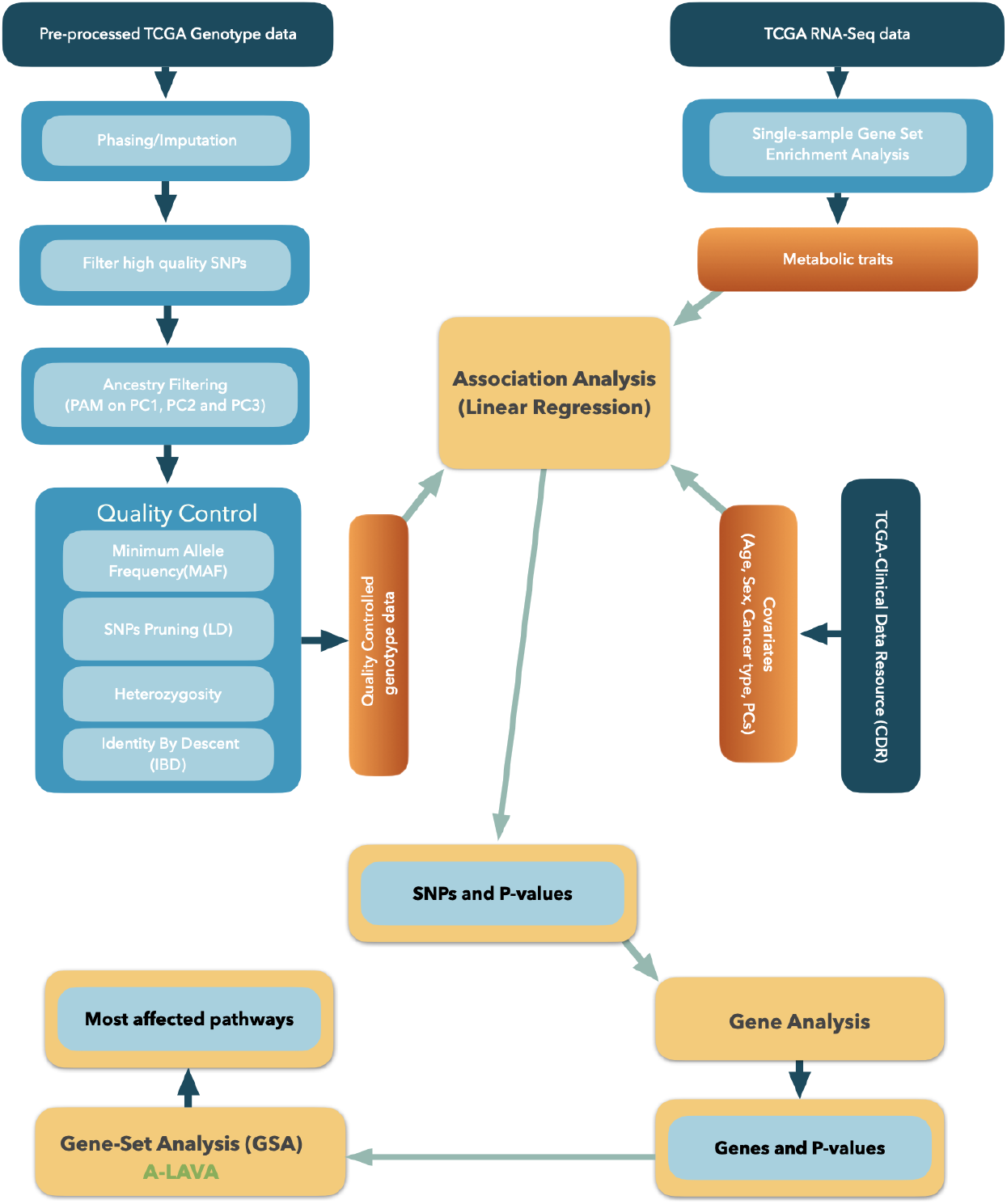
**A-LAVA overview. A genome-wide association study (GWAS) was conducted using TCGA germline SNP data. Samples were clustered by ancestry and underwent quality control (top left). Metabolic traits were derived from RNA-Seq data and refined via pathway enrichment scoring (top right). SNPs were mapped to autosomal genes and filtered based on strict statistical thresholds. Finally, the pathway analysis was performed using gene set analysis (GSA), which corrected for gene set overlaps to identify relevant networks**.

In the second step, gene-level analysis was done by mapping SNPs to their corresponding autosomal genes and filtering significant associations based on predefined statistical thresholds. We applied stringent criteria to identify candidate and suggestive genes, ensuring that only biologically relevant associations were retained for further study.

Finally, pathway-level analysis was done with the gene set analysis (GSA) suite that was designed to correct for overlapping gene sets. A-LAVA incorporates binary indicators to account for shared genetic components across overlapping gene sets, thus refining significance assessments and ranking metabolic networks. This approach prevents confounded associations by ensuring adjustments across all gene sets, irrespective of the number of shared genes. The final output of GSA is the list of the most affected metabolic pathways.

### 2.1 Data Processing

#### 2.1.1 Genotypes

The TCGA dataset includes germline genotype data obtained from SNP arrays for 11,440 samples from 10,776 participants across 33 cancer types, sourced from the Genomic Data Commons (GDC) legacy archive [24]. For genotype preparation and analysis, we followed the protocol established by Sayaman et al. [48]. Array-based variant calls were downloaded from GDC. Quality control measures excluded SNPs and individuals with more than 5% missing data, excessive heterozygosity, and palindromic SNPs. Genotype imputation was performed using the Haplotype Reference Consortium (HRC) reference panel [33], with phasing carried out via Eagle v2.4 and imputation conducted through Minimac4 on the Michigan Imputation Server [8]. High-confidence variants were retained based on imputation quality scores (R2 *<* 0.5) and processed using PLINK 1.9 [44].

To account for population stratification, samples were clustered into four ancestry groups: Europeans (8,337), Asians (633), Africans (928), and Native Americans (228), following genetic clusters identified in Sayaman et al. [48]. Minor allele frequency (MAF) correction was applied, removing SNPs with MAF *<* 0.005. Linkage disequilibrium (LD) pruning filtered out highly correlated SNPs (*r* ^2^ *>* 0.25) using a window size of 200 variants, as recommended by Choi et al. [7]. Additional quality control measures removed individuals with extreme heterozygosity (*>* 3 standard deviations from the mean per ancestry cluster) and excluded closely related individuals 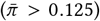 based on identity-by-descent (IBD) analysis [44]. We found that these strategies and the associated parameters help us minimize redundancy and reduce computational burden in the later steps. Note, however, that our choices might not be optimal in general and that alternative pre-processing strategies (such as LD clumping and so on) have been recently reported [2, 25].

After filtering, 8,579,009 SNPs were retained for 8,155 unrelated European individuals, 15,571,100 SNPs for 909 Africans, 7,358,496 SNPs for 610 Asians, and 9,570,879 SNPs for 221 Native Americans. Covariates such as age, sex, cancer type, and the top nine (9) principal components (PCs) were included to control for confounding effects, with the top nine PCs selected to capture 90% of the variance. Age was standardized, and cancer types with fewer than 100 cases were excluded for statistical stability. Covariate data were formatted for PLINK 2 [6] for downstream analysis.

#### 2.1.2 Metabolic pathways as target traits

TCGA RNA-Seq data provides insights into cellular transcriptomes by identifying activated genes and their transcription levels. To measure metabolic traits for association analysis, RNA-Seq data were processed to determine enrichment levels of predefined gene networks. Using single-sample gene set enrichment analysis (ssGSEA), enrichment scores were obtained for metabolic pathways and incorporated as target phenotypes in GWAS to investigate germline variants linked to these pathways. From this point forward, we refer to these pathways as “metabolic traits” to clarify their role as target traits in the GWAS analysis. Note that RNA-seq was not used for variant calling in the previous step.

TCGA RNA-seq expression data, encompassing 20,531 genes and approximately 11,000 samples, were sourced from the NIH Genomics Data Commons (GDC) archive ^1^. Gene set enrichment scores were computed using the ssGSEA method from the R GSVA package [21], with reference annotations (c2BroadSets) in R 4.2.1 [55]. Scores were filtered to retain 65 KEGG metabolic pathways. The KEGG pathways used in the association analysis and their mean enrichment scores are summarized in Supplementary Figure 21. Final gene set enrichment scores were formatted for PLINK 2 [6] and used as target metabolic traits in GWAS.

### 2.2 SNP and Gene-level Analysis

SNP-level association analysis was conducted using PLINK 2 [6], evaluating metabolic traits across 65 pathways. Genotype coding assigned 0 to homozygous reference alleles, 1 to heterozygous, and 2 to homozygous alternative alleles. Linear regression models assessed associations between allele counts and metabolic traits, producing *p*-values and effect sizes. Bonferroni-corrected significance thresholds were set at *p <* 5.8 × 10^−9^ for European populations, with population-specific thresholds of *p <* 5.2 × 10^−9^, *p <* 3.2 × 10^−9^, and *p <* 6.8 × 10^−9^ for Native American, African, and Asian groups, respectively. SNPs were annotated using dbSNP release 153 [52] and the Ensembl Variant Effect Predictor (VEP) [39], aligned to genome assembly GRCh37 (hg19).

To mitigate false positives, multiple testing correction applied the false discovery rate (FDR) [1], retaining SNPs with *p <* 0.05 for further analysis. Clumping was performed in PLINK 1.9 [43] to remove redundant correlated variants, reducing SNPs per locus to a single representative variant. While we used the PLINK’s default *p*-value settings in this analysis, we note that the more stringent *p*-value thresholds have been recommended in the literature [26, 60].

Gene-level analysis in MAGMA mapped SNPs to 18,114 genes using European reference data and computed gene-level *p*-values. Significant genes were identified using a Bonferroni-corrected threshold of 2.8 × 10^−6^ [62], with suggestive genes selected at *p <* 2.9 × 10^−5^ [50]. Gene mappings were refined using Entrez IDs [35] and converted to HGNC symbols via the R biomaRt package [16]. These processed SNP-level and gene-level results informed subsequent pathway analysis.

### 2.3 A-LAVA’s GSA Model

After identifying significant SNPs and genes, we sought to determine the most significant pathways or gene sets. In this study, the terms *pathways* and *gene sets* are used interchangeably. Both refer to sets of genes grouped based on shared functions, as defined in KEGG. Gene set analysis in MAGMA [9] employs *competitive gene set analysis*, testing whether genes in a given set show stronger associations with the phenotype than other genes. This approach evaluates a null hypothesis stating that no gene within the set is more strongly associated with the phenotype than those outside. Note that we here only directly compare against MAGMA; this is because A-LAVA and MAGMA share a common base and hence allow for an unbiased comparison, and because MAGMA is established as the canonical GSA tool for GWAS-related pipelines [9, 29, 49].

The model incorporates all genes in the dataset within a regression framework, defining a binary indicator variable *S*_*s*_ with elements *s*_*g*_, where *s*_*g*_ = 1 for genes within the set and *s*_*g*_ = 0 otherwise. Incorporating *S*_*s*_ as a predictor in the regression yields:

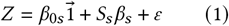

The parameter *β*_*s*_ quantifies the difference in association between gene set members and non-members. Testing the null hypothesis *β*_*s*_ = 0 against the one-sided alternative *β*_*s*_ *>* 0 provides a competitive assessment for each gene set, conceptually equivalent to a one-sided two-sample t-test comparing mean associations [10].

Gene-level analysis begins by calculating the association between genotype data and phenotypes, producing a matrix where each row corresponds to a gene. At the gene set level, a binary indicator variable is introduced, marking genes within the set as 1 and others as 0. Gene set analysis follows a bivariate framework, using genes as analysis units to determine whether the collective association of genes within the set exceeds that of genes outside the set (competitive analysis).

We improved the pipeline for identifying the most significant pathways using a gene-level multivariate regression model. ALAVA’s approach defines a separate binary indicator for all the gene sets and includes them in a regression model simultaneously, which transforms the above model (1) to:

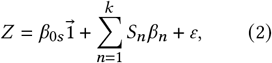

where *β*_0*s*_ is the regression constant term, and *k* denotes the number of gene sets included in the gene set analysis (GSA). Figure 2 illustrates how shared genes across gene sets 1 to *k* influence regression outcomes when testing *β*_*s*_ = 0 versus the one-sided alternative *β*_*s*_ *>* 0.

**Figure 2:**
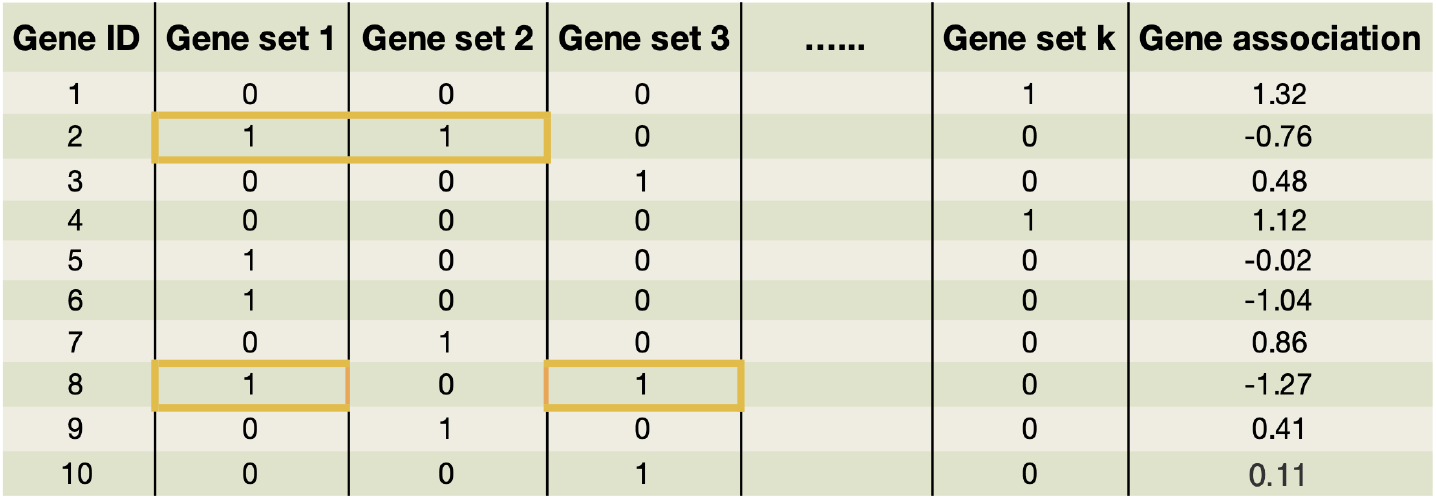
**Gene set competitive analysis in A-LAVA. This example is composed of 10 genes spread among** *k* **gene sets with their corresponding predictor of** *Z* **(gene association score). There is a separate binary indicator for each gene set (from 1 to** *k***). The orange rectangles show where there are shared genes between gene sets. In MAGMA, each binary indicator is entered separately into the linear regression model without taking into account the potential confounding effects arising from shared genes between gene sets. On the other hand, A-LAVA takes this information into account. Here, A-LAVA incorporates the fact that gene 2 is shared between gene sets 1 and 2, and that gene 8 is shared between gene sets 1 and 3**.

MAGMA’s competitive gene set analysis uses a generalized model that performs a conditional test of *β*_*s*_, adjusting for gene size and gene density. However, it does not account for genes shared across multiple gene sets. Additionally, MAGMA applies separate linear regression models for each gene set to evaluate significance. In contrast, A-LAVA simultaneously examines all gene sets, incorporating shared genes to estimate their coefficients (*β*) and corresponding *p*-values. This approach enhances computational efficiency while correcting *β* and *p*-values, refining gene set rankings within the study.

## 3 RESULTS

### 3.1 A-LAVA Simulations

In order to simulate the effects of overlap correction done by A-LAVA, we simulated *Z* as seen in formulas (1) and (2). We defined a vector of non-negative *Z* -scores for each gene where a larger *Z* -score indicates greater gene significance. The binary indicator vector *S*_𝓃_ follows gene sets from KEGG, where *S*_1_, *S*_2_, … represent the indicators for each gene set. Each gene set furthermore shares gene overlap with others, leading to the concept of a focus gene set—one that overlaps with all significant gene sets. Genes in significant gene sets were assigned *Z* -scores from a uniform distribution within [2.5, 3.0], whereas insignificant genes received scores within [0.0, 1.5]. A seed value of 5 ensured reproducibility. To quantify the significance within a focus gene set, we define the *ratio of significant genes* (RSG):

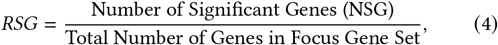

where the total number of genes corresponds to the sum of binary indicators within the gene set. The target files with the simulated *Z* - scores were then processed with A-LAVA and MAGMA. The output files were sorted according to t-value. A positive *β* and a *p*-value *<* 0.05 was the criteria for a significant gene set. This process was repeated such that each of the 65 KEGG gene sets became the focus gene set; hence, a total of 65 simulations were performed.

A-LAVA effectively refines pathway significance assessment by correcting for gene overlap, improving accuracy compared to MAGMA. MAGMA identifies a higher number of focus metabolic pathways as significant, largely due to unadjusted overlap effects (Figure 3). In contrast, A-LAVA applies corrections for shared genetic components, ensuring that only biologically relevant pathways are retained. This resulted in higher precision and *F*_1_ score (Figure 3), as well as consistently lower *β*-values, thus implying the removal of confounding effects.

**Figure 3:**
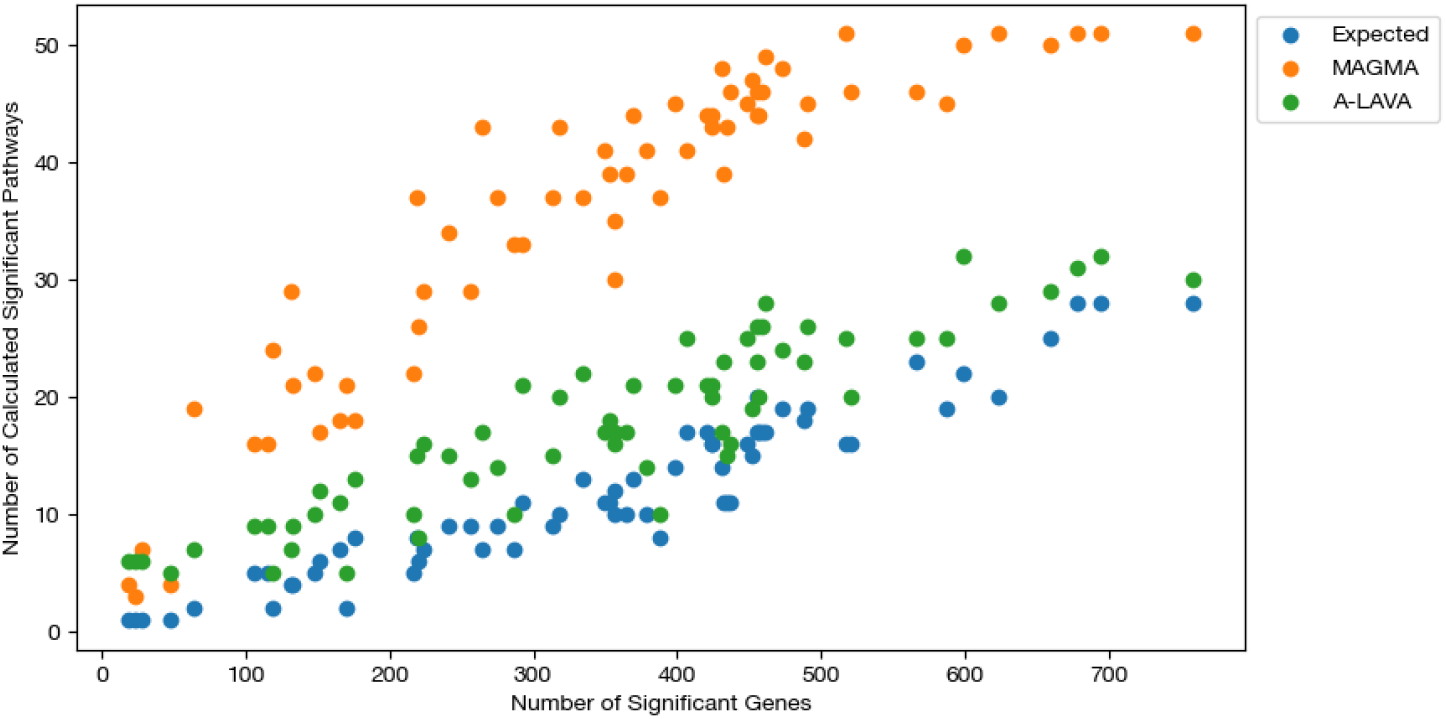
**Number of significant metabolic gene sets identified by A-LAVA and MAGMA across varying ranges of significant genes in simulation. Expected results are indicated in blue. MAGMA (orange) reports more spurious pathways due to uncorrected gene set overlap, which inflates significance. In contrast, A-LAVA (green) adjusts for overlapping genes, yielding more reliable associations, thus highlighting its effectiveness in identifying robust metabolic signals**.

The only observed downside is the slight decrease in recall due to few false negatives, which is suspected to have caused a single unexpected case with an RSG of 1.0 that was found insignificant under A-LAVA. This implies a potential over-correction of gene overlap correction in certain rare cases.

Figure 4 highlights key differences between MAGMA and A-LAVA in significance assessment. A-LAVA correctly identifies more true negative (TN) scenarios, ensuring non-significant pathways remain insignificant, while MAGMA frequently misclassifies originally insignificant pathways due to overlap (false positives). Only A-LAVA detects cases where overlapping pathways remain insignificant. Across 65 simulations, A-LAVA consistently corrected for gene overlap more effectively than MAGMA, thus reducing the number of false positives. Overall, this demonstrates that A-LAVA is able to provide more precise pathway rankings and improved statistical reliability in GSA.

**Figure 4:**
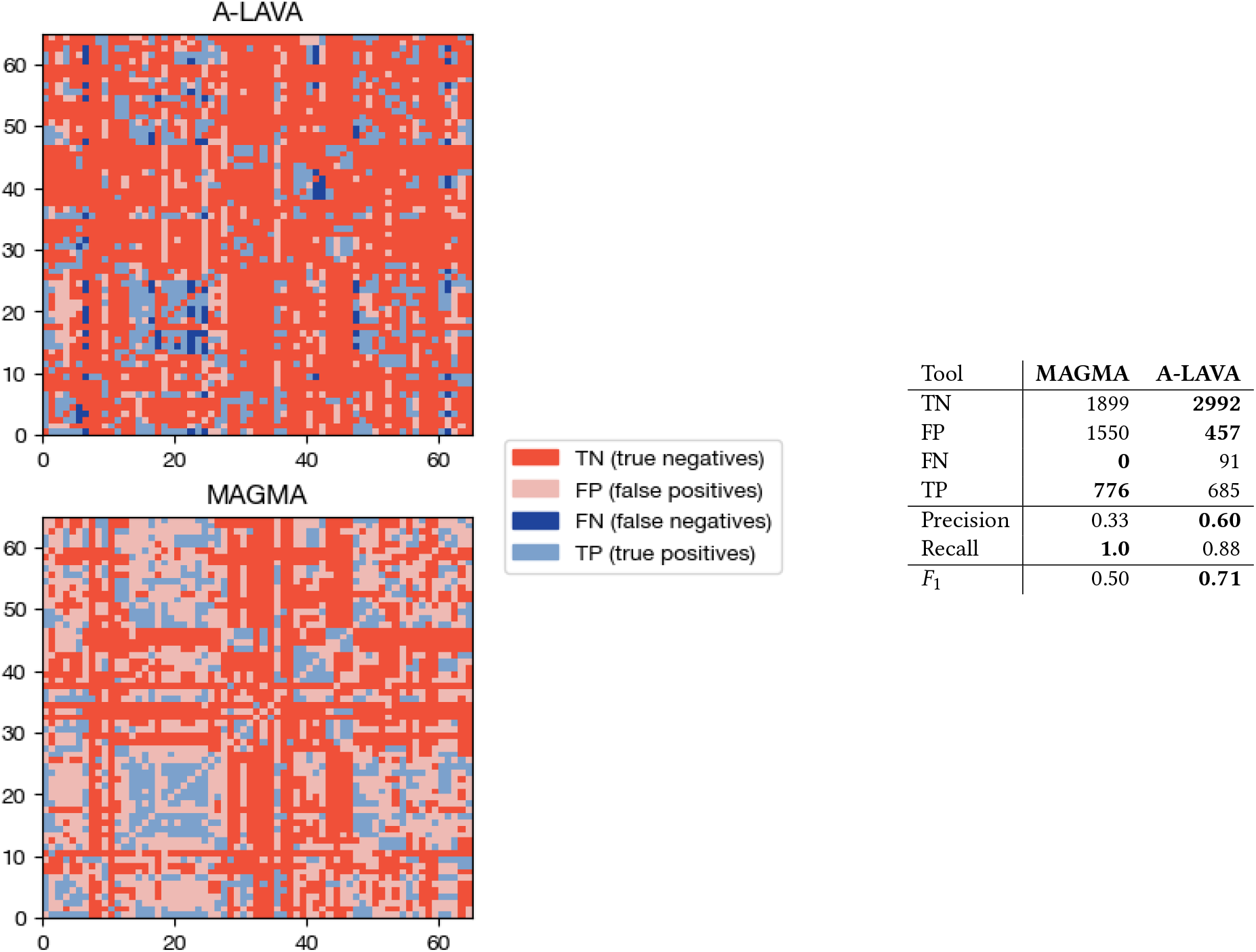
**Significance assessment differences between MAGMA and A-LAVA on simulated dataset. (Left) Each row represents a focus gene set, and each column represents the 65 gene sets in the same order. The four colours refer to the results drawn by A-LAVA and MAGMA after selecting any (row, column) entry in the matrix. These results are based on the following attributes labelled for brevity: whether the focus gene set overlaps with the gene set in this column and whether the method found the gene set in this column significant. (Right) Summary statistics of the differences between A-LAVA and MAGMA in the tabular form. A-LAVA corrects large number of false positives (and significantly improves precision and** *F*_1_ **score) at the expense of a few false negatives (slightly lower recall). Bold text indicates better result**.

### 3.2 A-LAVA TCGA Results

#### 3.2.1 SNP and Gene-level Results

GWAS summary statistics were visualized using Manhattan and quantile-quantile (Q-Q) plots (Supplementary Figures 11 and 12), illustrating genomic association strength and *p*-value distribution. GWAS analyses for Asian, Native American, and African populations yielded excessive SNP associations within significance thresholds, with 6,699 SNPs identified in the Native American population despite a sample size of only 221. Genomic inflation factors (λ) deviated from the expected value of 1, indicating stratification biases (Supplementary Figures 23, 25, and 27) [56]. Given that large sample sizes are preferred in GWAS to minimize biases [17], these populations were excluded from further analysis due to extreme stratification effects and limited data availability.

For Europeans, 71 genome-wide significant SNPs were linked to 10 metabolic traits (*p <* 5.8 × 10^−9^), with glutathione metabolism showing the strongest associations (31 SNPs). Notably, rs36209093 mapped to *GSTM2* (*p* = 1.52 × 10^−39^), reinforcing its role in detoxification pathways [45]. LD clumping identified 20 independent loci across 10 traits, with 14 novel associations not previously reported in the GWAS catalogue (Supplementary Figure 13) ^2^, thus highlighting the importance of correcting for population stratification and refining association signals for enhanced metabolic trait discovery.

All SNPs were mapped to 18,114 autosomal genes, with significance determined using a Bonferroni-corrected threshold of 2.8 × 10^−6^ and suggestive significance set at 2.9 × 10^−5^. We identified 19 candidate genes associated with 16 metabolic traits and 30 suggestive genes linked to 23 metabolic traits. Glutathione metabolism and alpha-linolenic acid metabolism exhibited the most connections, with eight and five associations, respectively (Supplementary Figure 14). Among non-metabolic genes, *OR12D3, ZNF215*, and *SMLR1* showed the highest frequency of associations (Supplementary Figure 7). For metabolic genes, *GRHPR, SCN3A, GSTM1*, and *GSTM5* were frequently detected, with the glutathione S-transferase (GSTs) super-family (*GSTM1*–*GSTM5* and *GSTZ1*) appearing in nine instances (Supplementary Figure 7).

Biologically significant associations include *LIMD1* and *VARS2*, linked to valine leucine and isoleucine biosynthesis, both exhibiting negative effect sizes. Downregulation of *VARS2* supports a metabolic phenotype characteristic of cancer (Warburg effect [32]). Mutations in the GST super-family impair detoxification pathways, potentially affecting drug efficacy and cancer susceptibility [45]. Elevated GSTP expression in tumour cells suggests mechanisms beyond drug detoxification, reinforcing the need to consider GSTM variants in cancer treatment strategies. Other identified genes include *FADS2*, associated with alpha-linolenic acid metabolism, and *DHFR*, involved in one-carbon metabolism and folate biosynthesis. Detection of *DHFR* aligns with its known role in folate processing, converting dietary folate to THF, a critical one-carbon acceptor. Additionally, *MSH3* was linked to DNA repair and base excision repair pathways, known to influence cancer progression [23, 31]. Overall, these findings highlight the key metabolic genes and their relevance in disease susceptibility.

#### 3.2.2 GSA analysis

We used A-LAVA’s GSA model to analyze the pathway selection and compared it to MAGMA across 65 metabolic traits. As shown in Figure 5, the drug metabolism Cytochrome P450 trait revealed significant shifts in *p*-values and *β*-values. For example, MAGMA ranked `hsa00980` as the top pathway, but A-LAVA replaced it with `hsa00260` while adjusting its *p*-value from 0.00007 to 0.02059. Additionally, several pathways were removed following overlap correction. Using a threshold of *p <* 0.05, only five pathways remained significant under A-LAVA, compared to 16 in MAGMA, further highlighting the necessity of correcting for shared genes.

**Figure 5:**
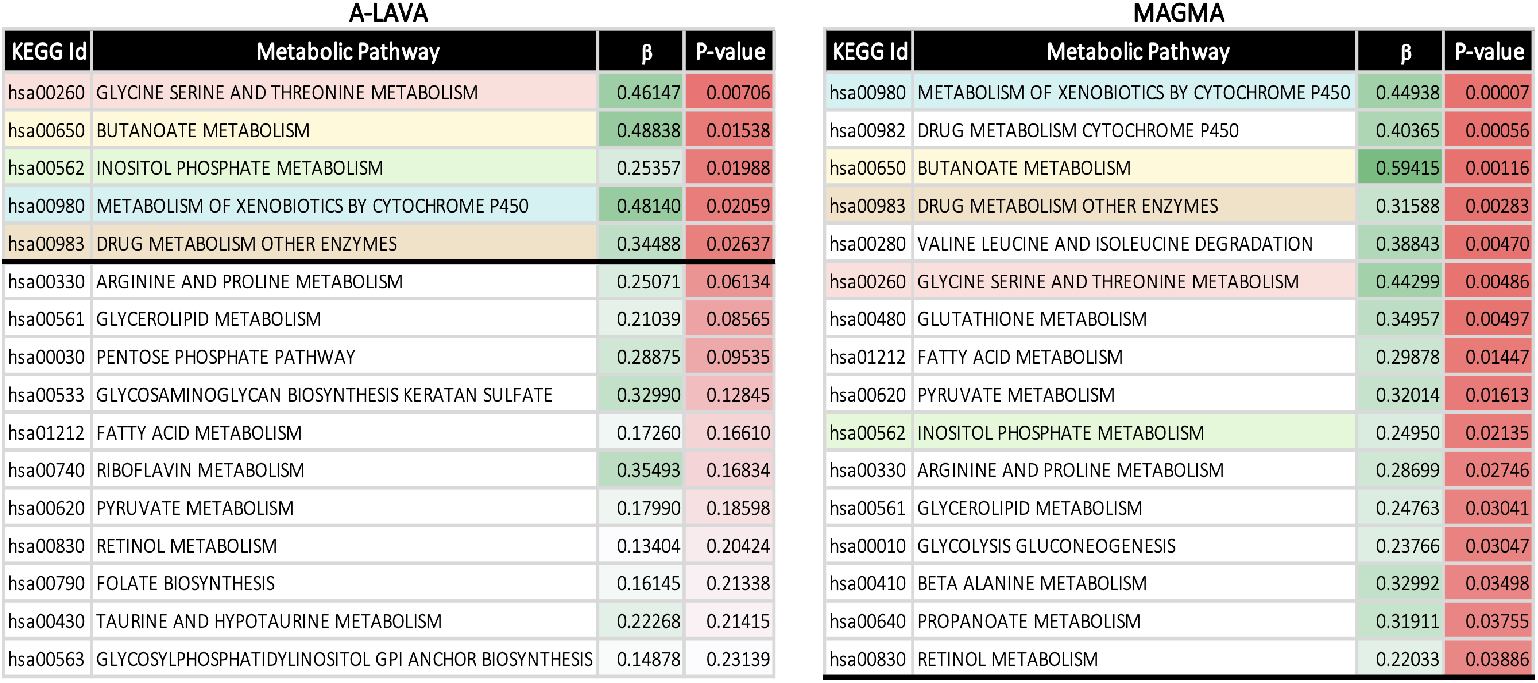
**The top gene sets selected using the A-LAVA (left) and MAGMA (right) are shown for a key drug metabolism trait (cytochrome P450). The top gene sets in common between both methods have the same colour shade on both sides. *P* -values and** *β* **values are represented using red and green colours to highlight the outcome discrepancy between the two methods. The bold black line is where the cut-off threshold meets the ranked gene set list. Here, A-LAVA selects fewer gene sets and puts them in a different order as compared to MAGMA**.

Figure 6 illustrates the correlation shifts induced by A-LAVA, reinforcing the importance of adjusting for confounding genetic overlap. Across all traits analyzed, A-LAVA consistently refined rankings, improving accuracy in identifying truly significant pathways.

**Figure 6:**
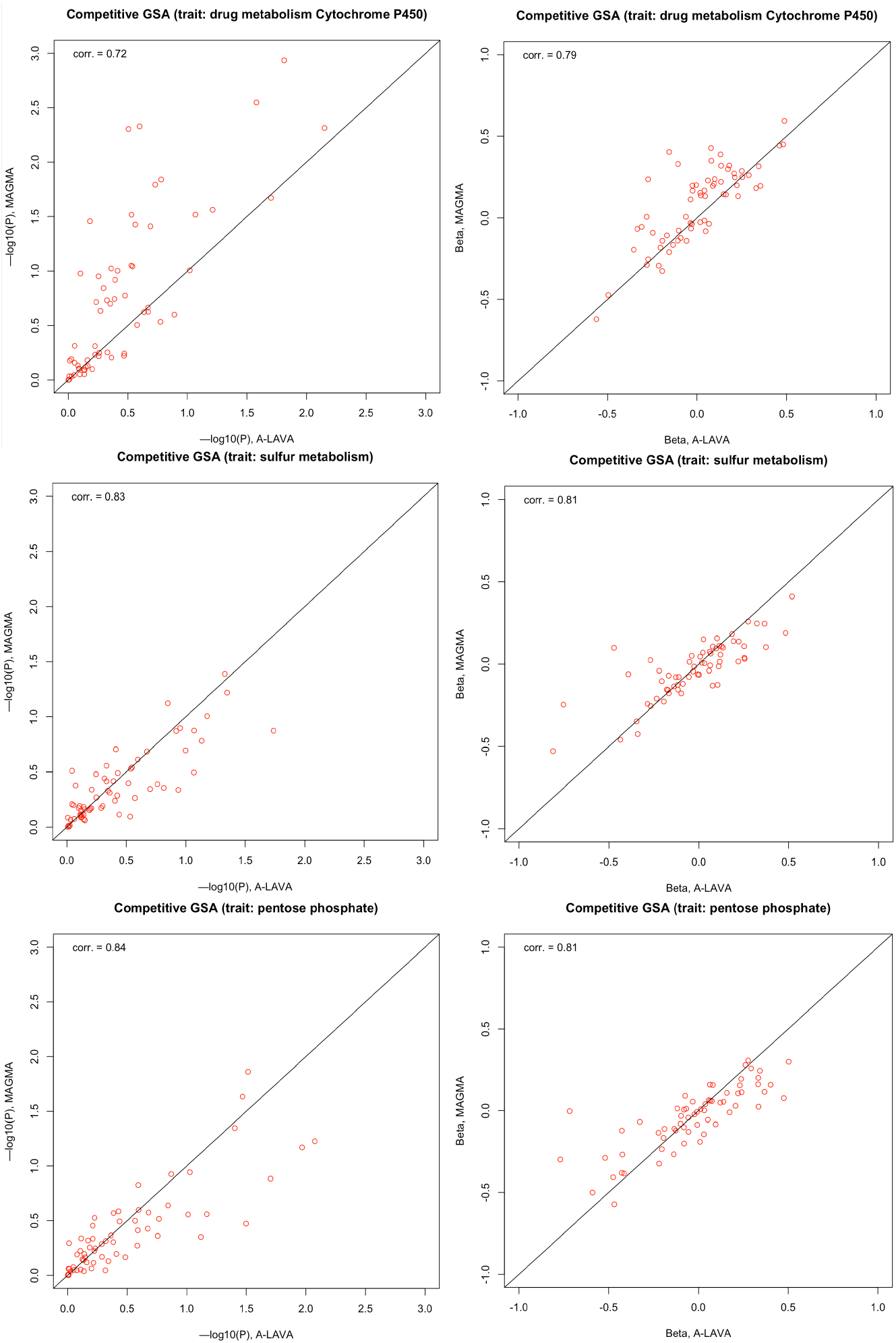
**Pearson correlation coefficient illustrating the similarity of outcomes in A-LAVA (*x*-axis) and MAGMA (*y*-axis) for a subset of studied metabolic traits. The plots depict a significant shift in** *p***-values (left) and** *β* **(right) resulting from correcting for shared genes, thus showing that the shared genes are indeed a confounding factor that should be addressed**.

Finally, Supplementary Figures 16 and 17 summarize frequency counts and statistical parameters for top metabolic pathways across all traits. Ether lipid metabolism emerged as the most frequently selected significant pathway, appearing in 28% of analyzed traits. Mean *p* and *β*-values further confirm the refined significance rankings, with pathways like glycan degradation and metabolism of xenobiotics by cytochrome P450 showing notable enrichment adjustments under A-LAVA.

Note that in all cases, both A-LAVA and MAGMA were nearly instantaneous when it comes to execution speed, and neither of them required significant computing power or RAM for execution.

These findings collectively demonstrate the effectiveness of A-LAVA in pathway selection, correcting gene overlap biases, and improving statistical reliability in gene set analysis methods.

## 4 LIMITATIONS AND FUTURE WORK

One of the limitations of the current work is the lack of correction for hidden biases inherent in the input data. For example, our pipeline currently assumes that the TCGA germline variant calls are accurate and leaves it to the end-user to correct for technical artifacts or potential impact of somatic mutations to the germline calls [5, 41]. We also note that a particular cancer type might also be a source of bias. While we have performed a joint analysis of various cancer types in this study, we plan to conduct an independent analysis for different cancer types in the future.

Another major limitation of this study is its reliance on the particular data preprocessing and filtering choices (e.g., [7, 48]) that might themselves introduce certain biases. It remains to be seen whether our pipeline is robust to other data preprocessing choices (such as LD pruning; e.g., [2]) or not. We also plan to explore the exact reasons for false negatives that might occasionally occur when correcting for overlapping gene sets.

Regarding the biological validity of our results, we aim to further validate in the wet lab whether the identified SNPs and mutated genes are true causal variants influencing the enrichment of metabolic pathways by leveraging CRISPR-Cas9 technology to experimentally assess the impact of inherited variants on cellular metabolism. We also plan on exploring the effect of different *p*-value thresholds on the variant selection and their true metabolic effect.

## 5 CONCLUSION

Despite extensive efforts in cancer genomics, the role of germline variants in metabolic gene networks remains underexplored. We performed GWAS to uncover potential causal germline variations influencing metabolic traits across multiple cancer types and populations. Our findings revealed strong associations for glutathione metabolism, xenobiotics by cytochrome P450, and drug metabolism cytochrome P450 traits, with the glutathione S-transferase super-family emerging as the most significant gene group.

One key challenge in GWAS is detecting weak but genuine associations due to the large number of tests performed [59]. To overcome this, we employed gene set analysis, which aggregates signals across biologically related genes to provide deeper functional insights [9]. However, conventional gene set analysis methods consider important binary indicators independently and, as such, often suffer from bias due to overlapping gene sets, leading to confounded associations [11]. For that reason, our analysis suite A-LAVA effectively resolves these limitations by considering all such indicators at the same time to correct for gene overlap, ensuring precise pathway ranking. Unlike MAGMA, which does not adjust for shared genes, A-LAVA ensures that all overlapping gene sets are adjusted regardless of the number of shared genes, minimizes biases and improves statistical reliability. Our pathway-level analysis identified 201 significant pathways across 65 metabolic traits, reinforcing the hypothesis that correcting for overlapping confounding factors leads to more accurate *p* and *β*-values. Simulation results demonstrate A-LAVA’s impact, revealing significant shifts in pathway rankings, reducing false positives, and improving detection accuracy.

By addressing these biases, A-LAVA provides a refined framework for gene set analysis, improving genomic studies’ ability to uncover biologically meaningful metabolic trait associations. Its improved GSA modelling and detection capabilities enhance the prioritization of biologically relevant SNPs and genes, and offer valuable insights for researchers seeking to leverage metabolomics data in cancer prevention and treatment. It thus can guide future functional validation studies by refining predictions of population-specific susceptibility to treatments and medications, therefore ultimately advancing precision medicine strategies.

## A APPENDIX

**Figure 7:**
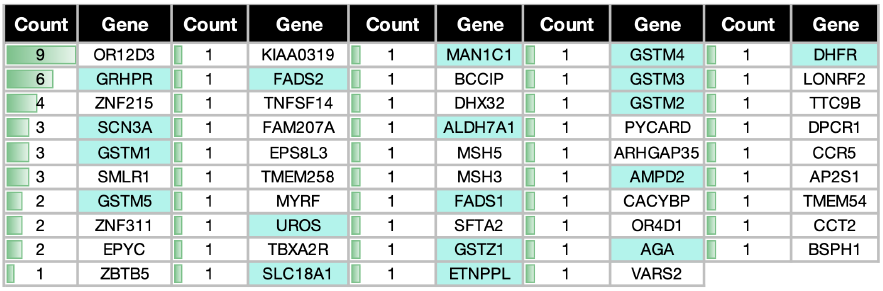
Ranking of genes based on the frequency of occurrence as top genes (MAGMA) among metabolic traits.

**Figure 8:**
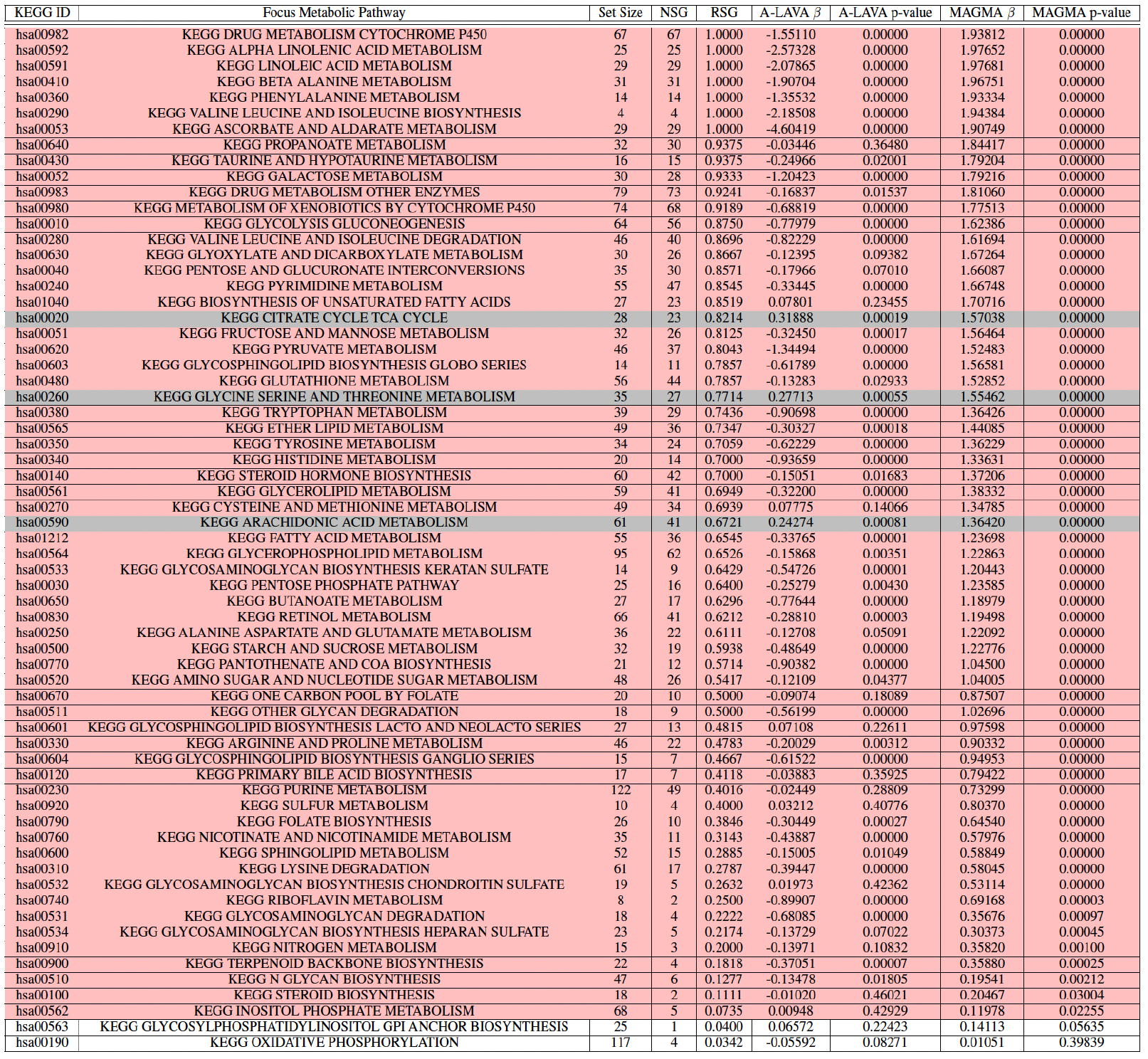
**The results of A-LAVA and MAGMA for the focus gene set for each simulation. The rows highlighted in pink are the focus gene sets that were found to be significant by only MAGMA. The rows highlighted in gray are the focus gene sets that were found to be significant by both A-LAVA and MAGMA. The rows that are not highlighted are the focus gene sets that were found to be insignificant by both A-LAVA and MAGMA. Gene sets are significant if they have a positive** *β* **value and a** *p***-value** *<* 0.05. **The chart is sorted by decreasing RSG value**.

**Figure 9:**
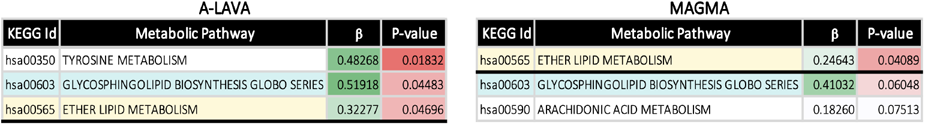
**The top gene sets selected using the A-LAVA (left) and MAGMA (right) for the sulfur metabolism trait. The top gene sets in common between both methods have the same colour shade**. *p* **and** *β* **values are represented using red and green colour values to highlight the outcome discrepancy between the two methods. The bold black line is where the cut-off threshold meets the ranked gene set list**.

**Figure 10:**
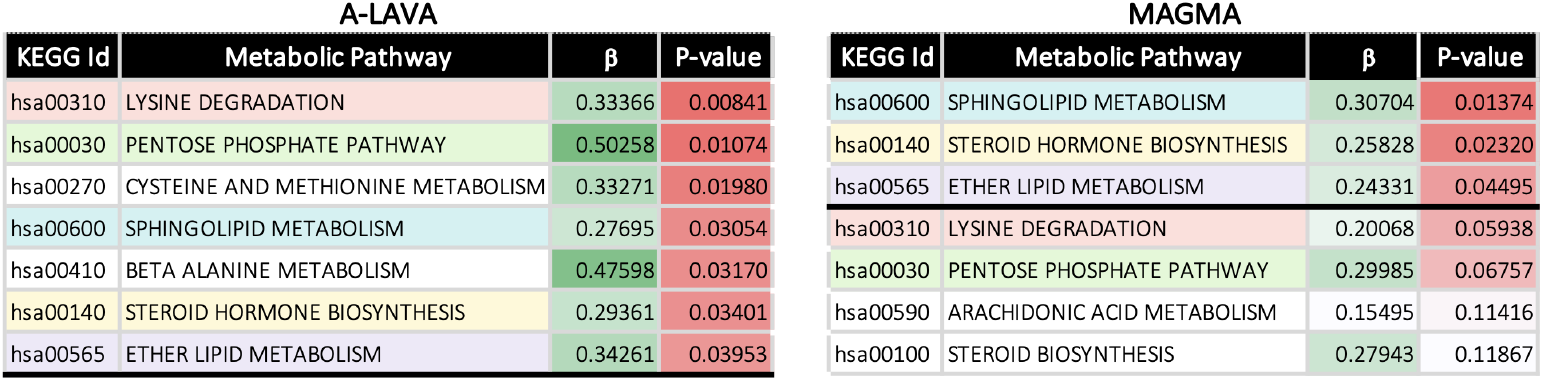
**The top gene sets selected using the A-LAVA (left) and MAGMA (right) for the pentose phosphate trait. The top gene sets in common between both methods have the same colour shade**. *p* **and** *β* **values are represented using red and green colour values to highlight the outcome discrepancy between the two methods. The bold black line is where the cut-off threshold meets the ranked gene set list**.

**Figure 11:**
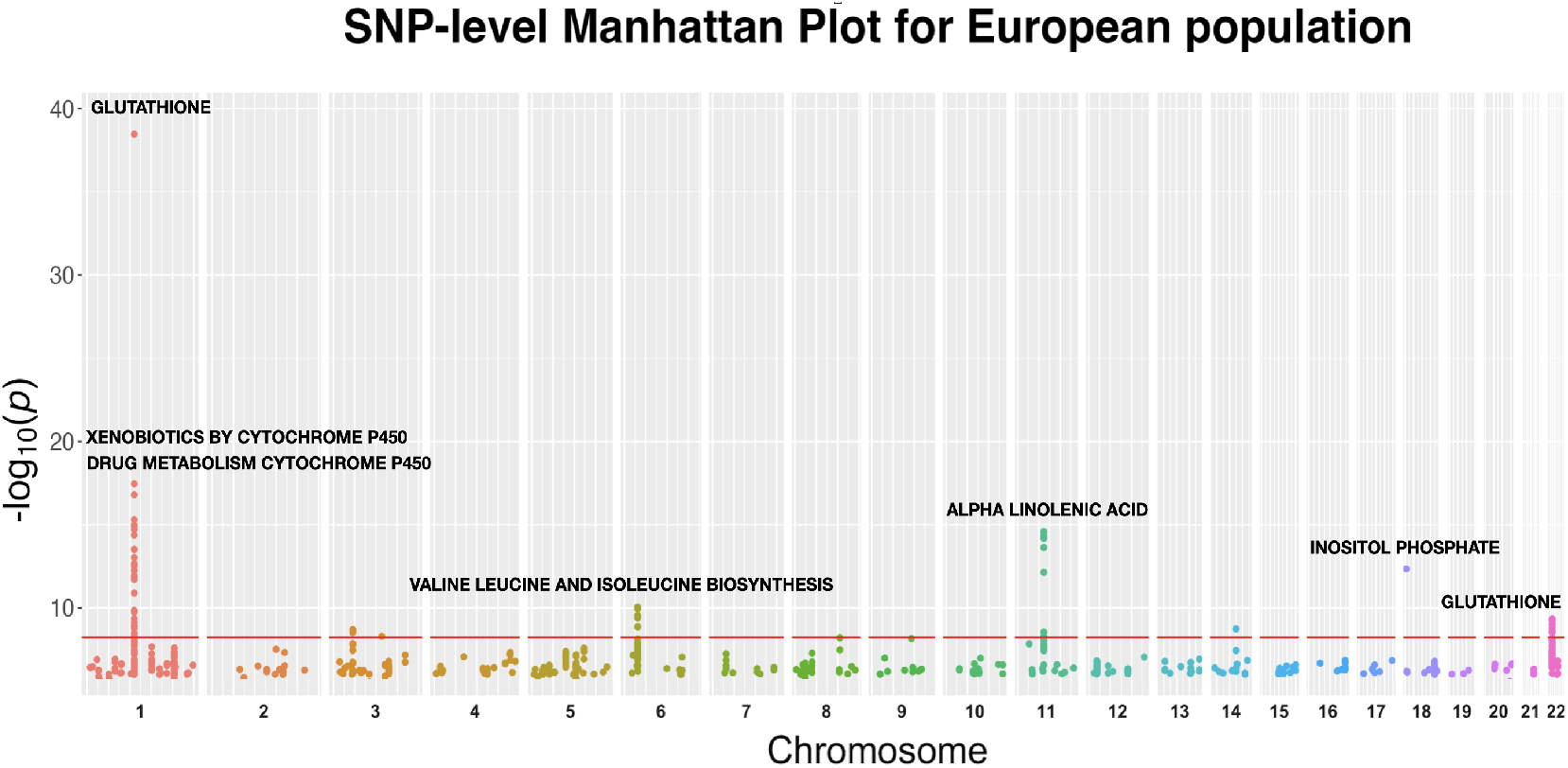
**SNP-level Manhattan plot showing all top SNPs across all traits in the European population. Each dot represents an SNP, with SNPs ordered on the** 𝓍 **-axis according to their genomic position**. *Y* **-axis represents the strength of their association measured as** − log_10_ **transformed** *p***-values starting from** 1 × 10^−6^. **The red line shows the threshold of genome-wide significance** (*p <* 5.8 × 10^−9^).

**Figure 12:**
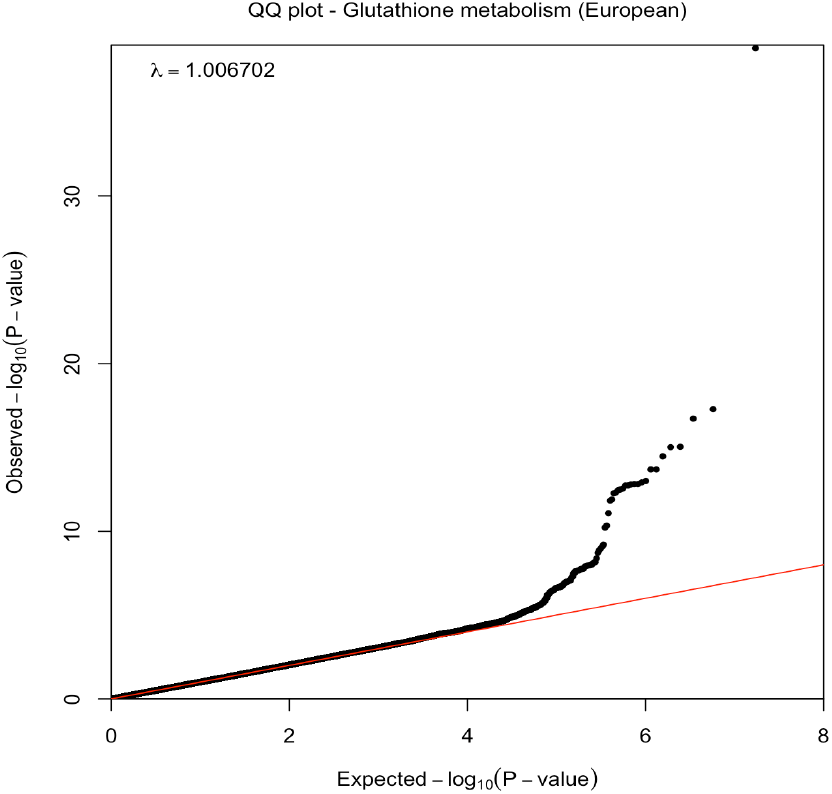
**Quantile–quantile plot illustrating the distribution of expected** *p***-values under a null model of no significance (red line) versus observed** *p***-values (black dots) for glutathione metabolism in the European population. The deviation of observed** *p***-values from the expected line highlights the significance of detected SNPs. The genomic inflation factor (**λ**) being close to 1 indicates no evidence of inflation, confirming the previously reported genetic homogeneity of the European samples [13]**.

**Figure 13:**
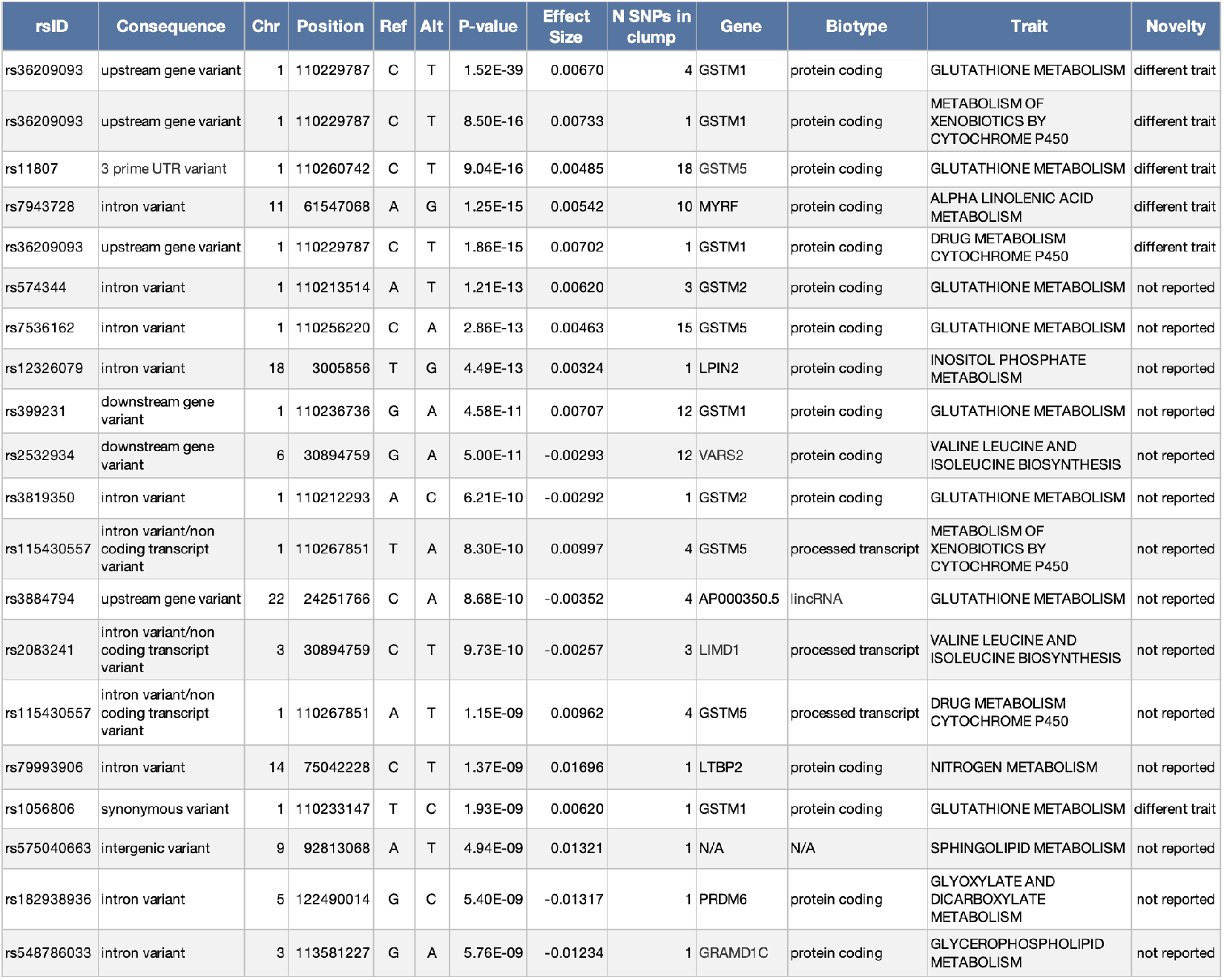
**The information of representative SNPs per region of LD associated with ten metabolic traits. The SNPs are annotated with rsIDs and are sorted based on their significance (increasing** *p***-value). The position of SNPs, the genes, and the transcript classification (biotype) they belong to, along with their consequence prediction, are also shown in this table. The effect size reflects the SNP’s contribution to the trait’s genetic variance, and the novelty column indicates whether the corresponding SNP has been previously reported**.

**Figure 14:**
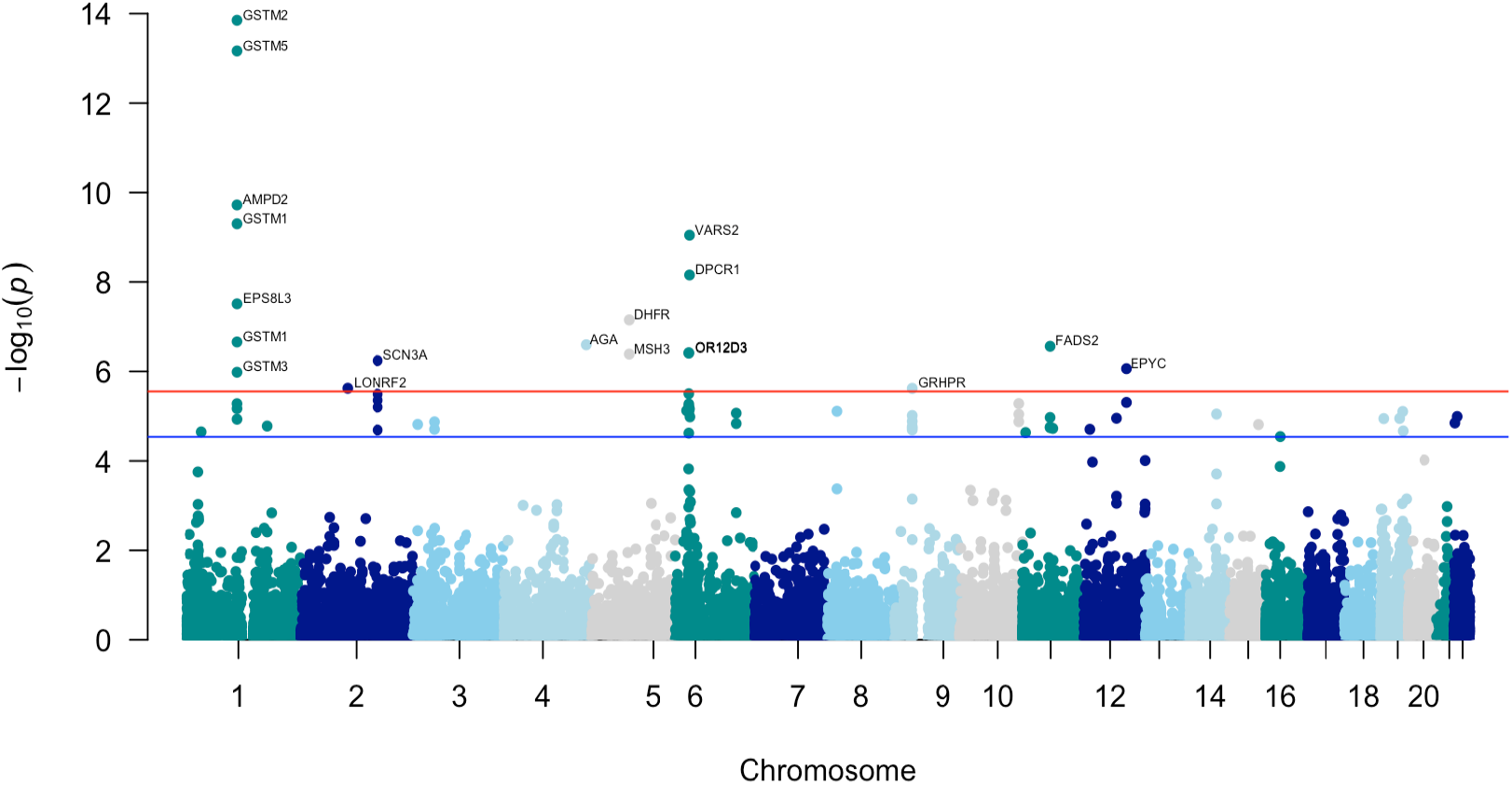
**Gene-level Manhattan plot where each point represents a gene. The red and blue lines correspond to the gene-level significance and suggestive thresholds**.

**Figure 15:**
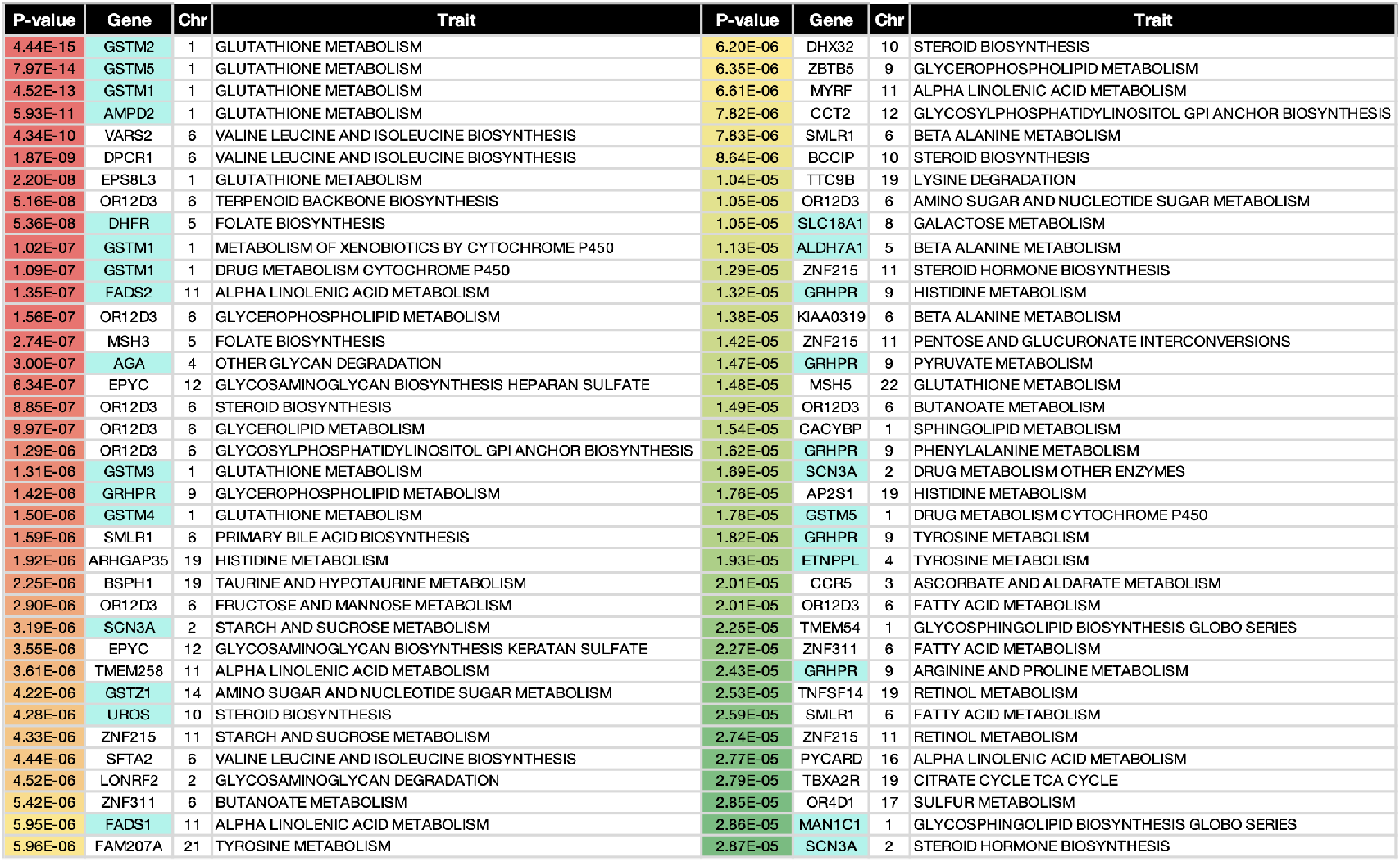
**List of top genes for European ethnic group**. *p***-values tend to be red and green, representing candidate and suggestive genes, respectively. The metabolic genes are highlighted in cyan blue, and the corresponding metabolic traits can be seen in the right-most columns**.

**Figure 16:**
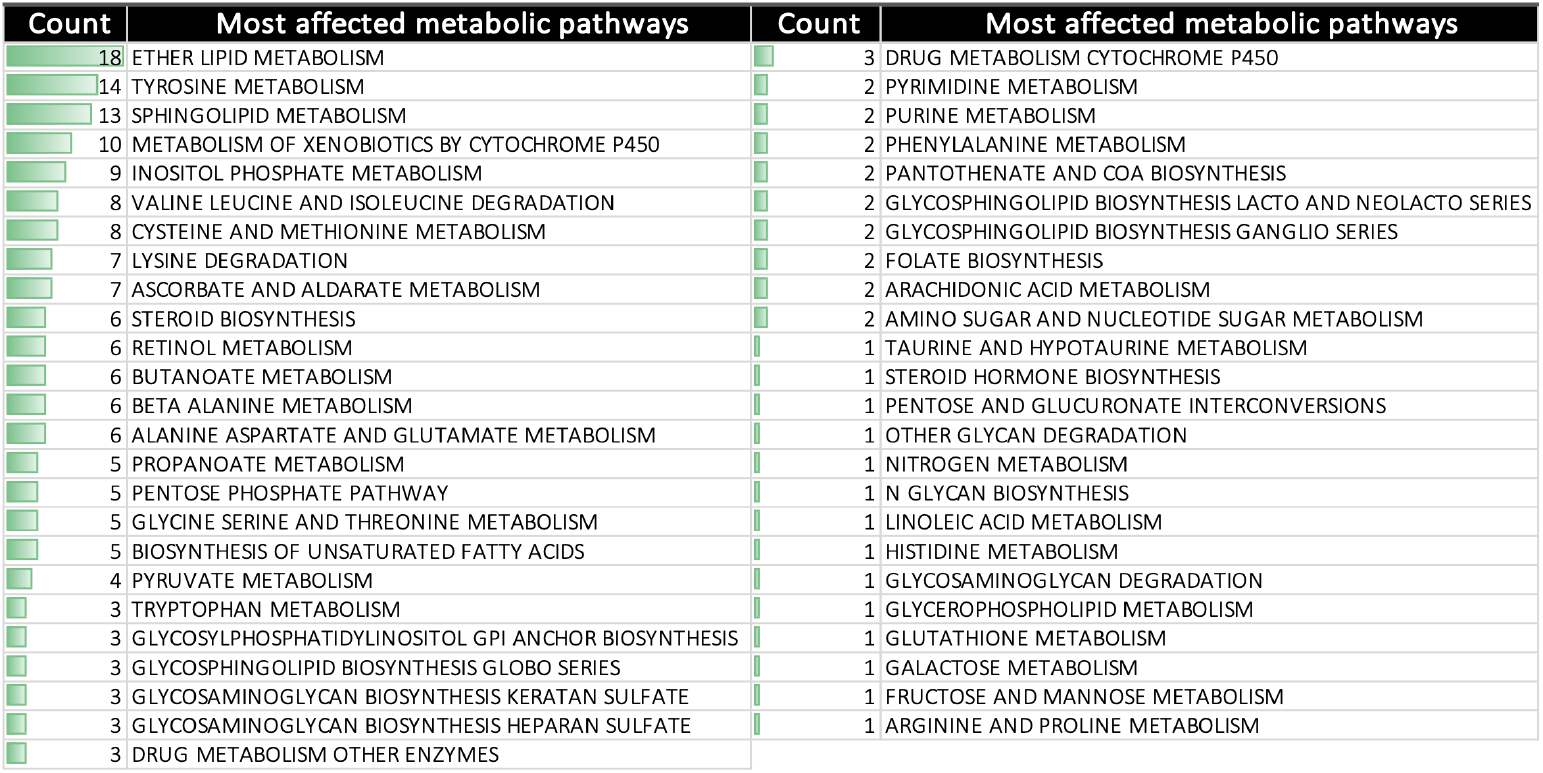
**List of the significant metabolic pathways and the count of their occurrence as significant in GSA across 65 traits using A-LAVA. The count measures are shown as green bars**.

**Figure 17:**
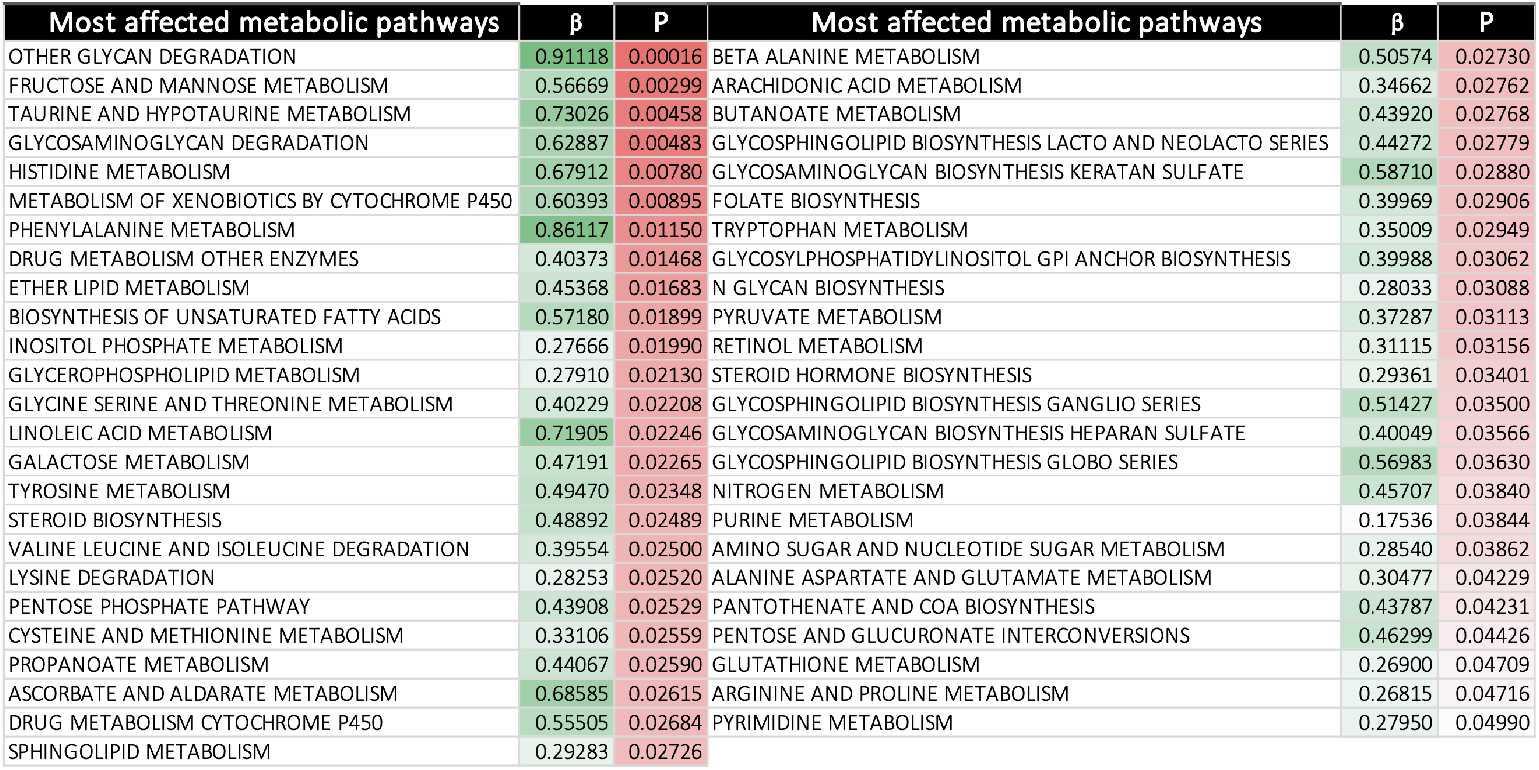
**Significant metabolic pathways and the mean of their** *p***-value and** *β* **across 65 traits. The pathways are listed in an increasing** *p***-value order, with darker red cells corresponding to lower** *p***-values. Also, the darker green cells highlight a more significant** *β* **value (difference in the association between genes in the pathway and genes outside the pathway)**.

**Figure 18:**
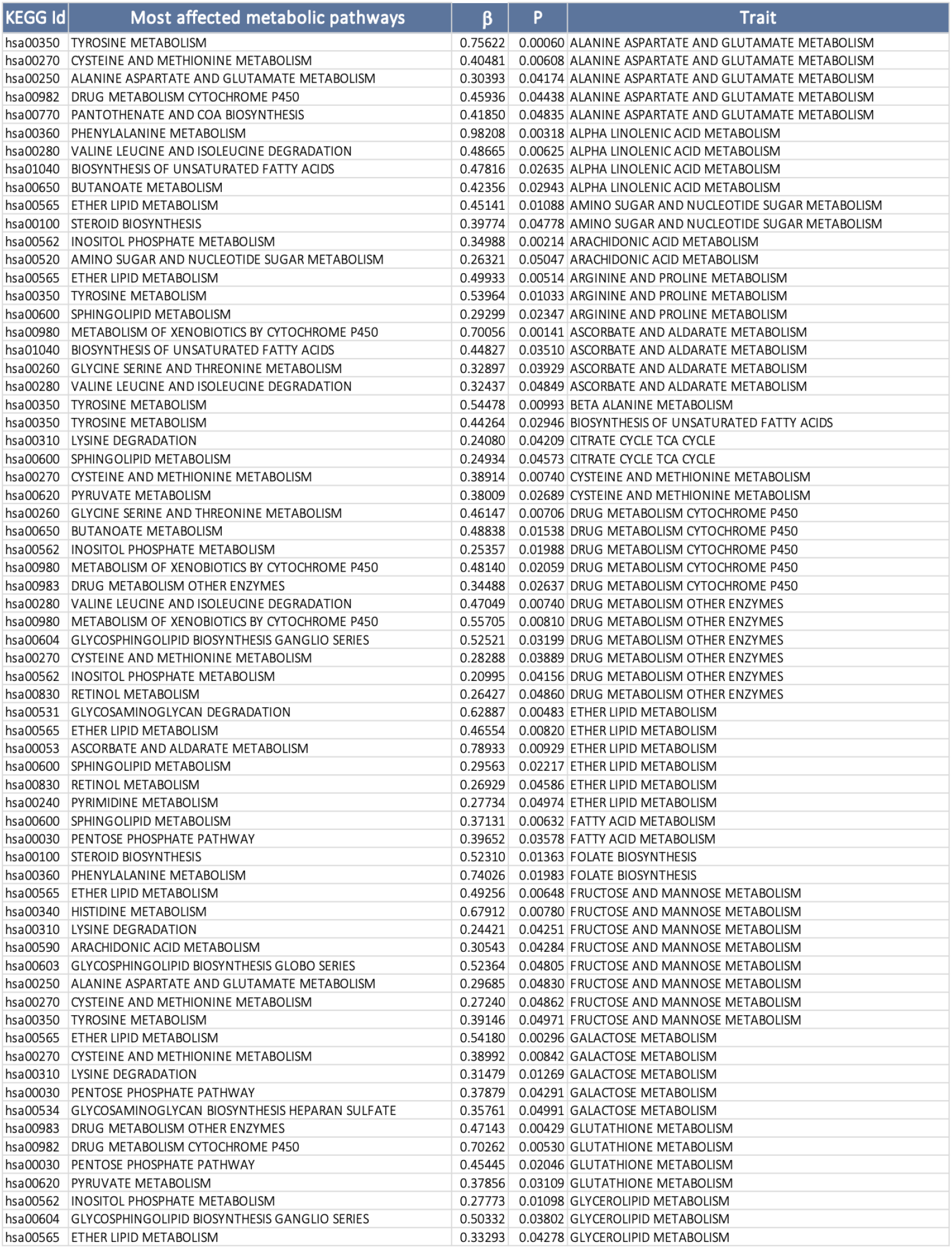
**Metabolic pathways that passed the GSA significance threshold in A-LAVA (part 1). The pathways are ranked in increasing** *p***-value order for each of the 65 initial traits. The corresponding** *β* **value suggests the difference in the association between genes included in the pathway and genes outside the pathway**.

**Figure 19:**
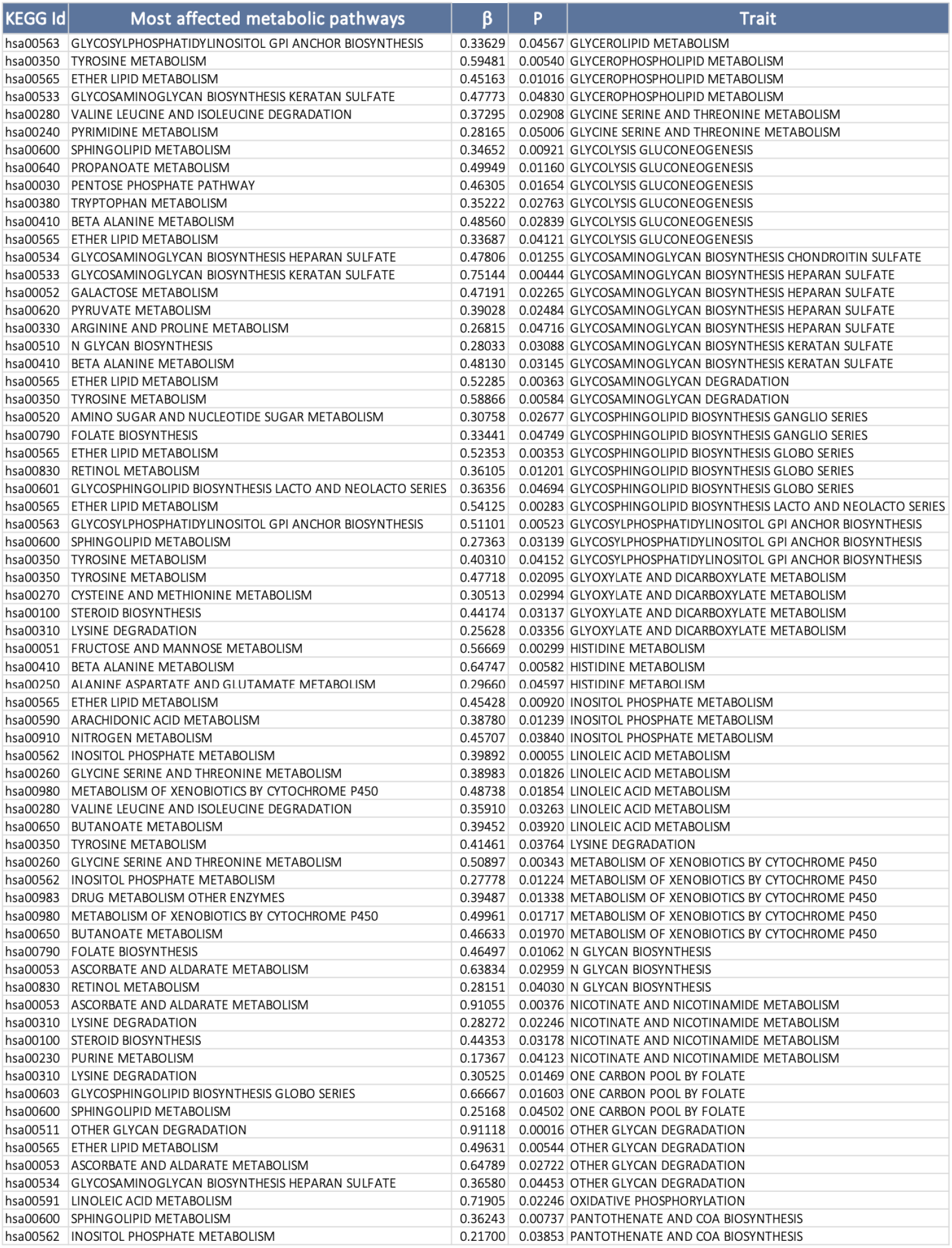
**Metabolic pathways that passed the GSA significance threshold in A-LAVA (part 2). The pathways are ranked in increasing** *p***-value order for each of the 65 initial traits. The corresponding** *β* **value suggests the difference in the association between genes included in the pathway and genes outside the pathway**.

**Figure 20:**
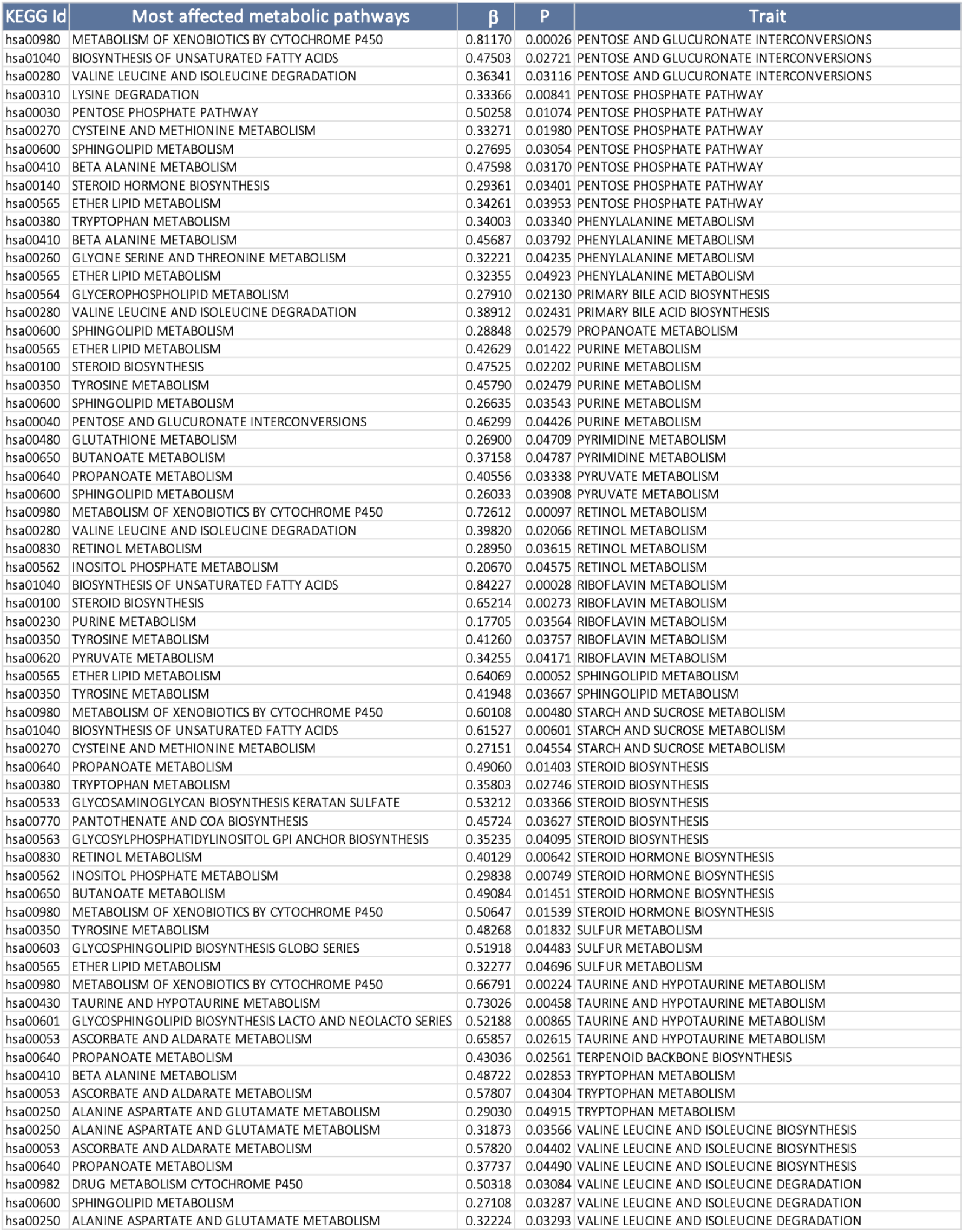
**Metabolic pathways that passed the GSA significance threshold in A-LAVA (part 3). The pathways are ranked in increasing** *p***-value order for each of the 65 initial traits. The corresponding** *β* **value suggests the difference in the association between genes included in the pathway and genes outside the pathway**.

**Figure 21:**
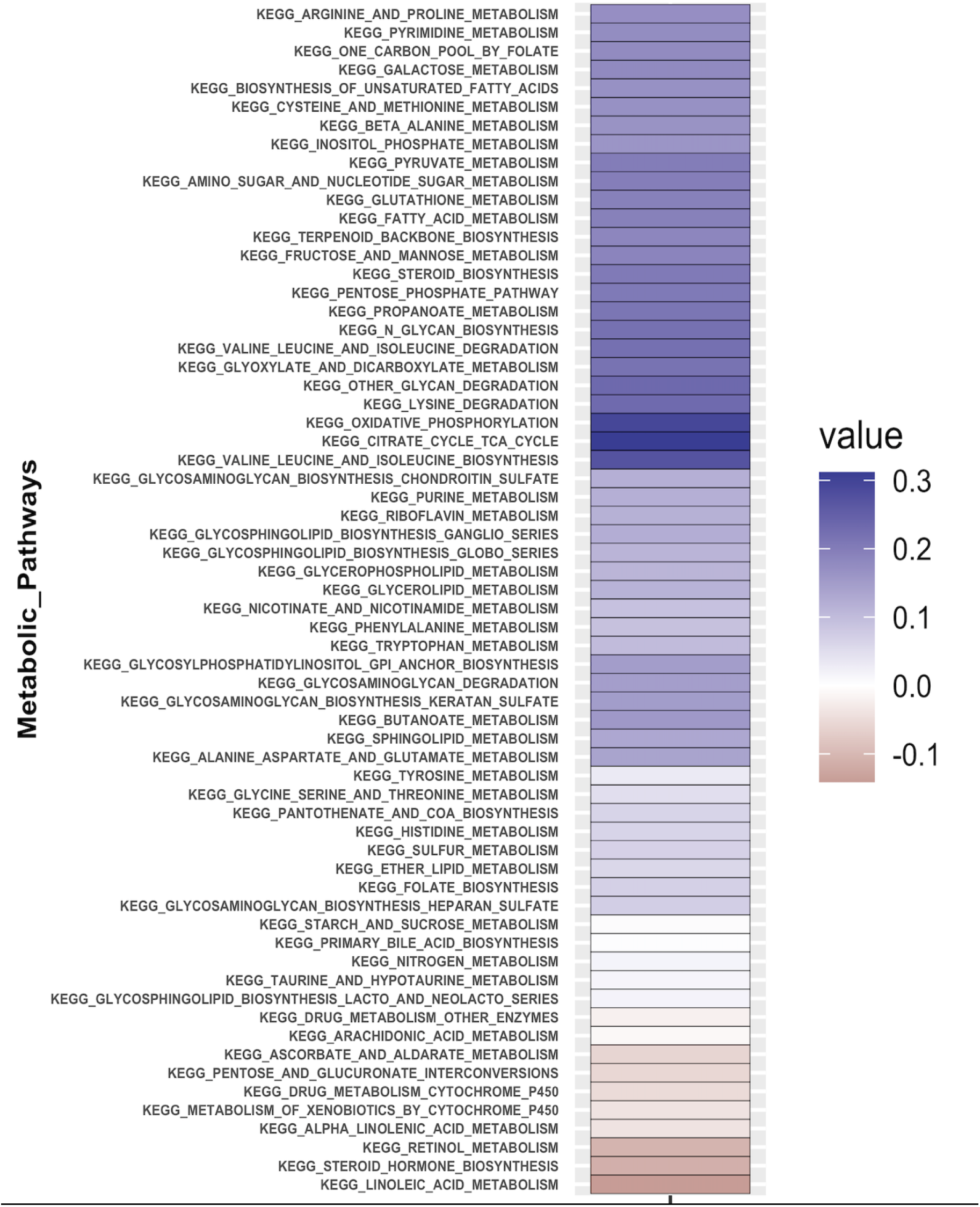
Metabolic pathways enrichment mean scores across all samples.

**Figure 22:**
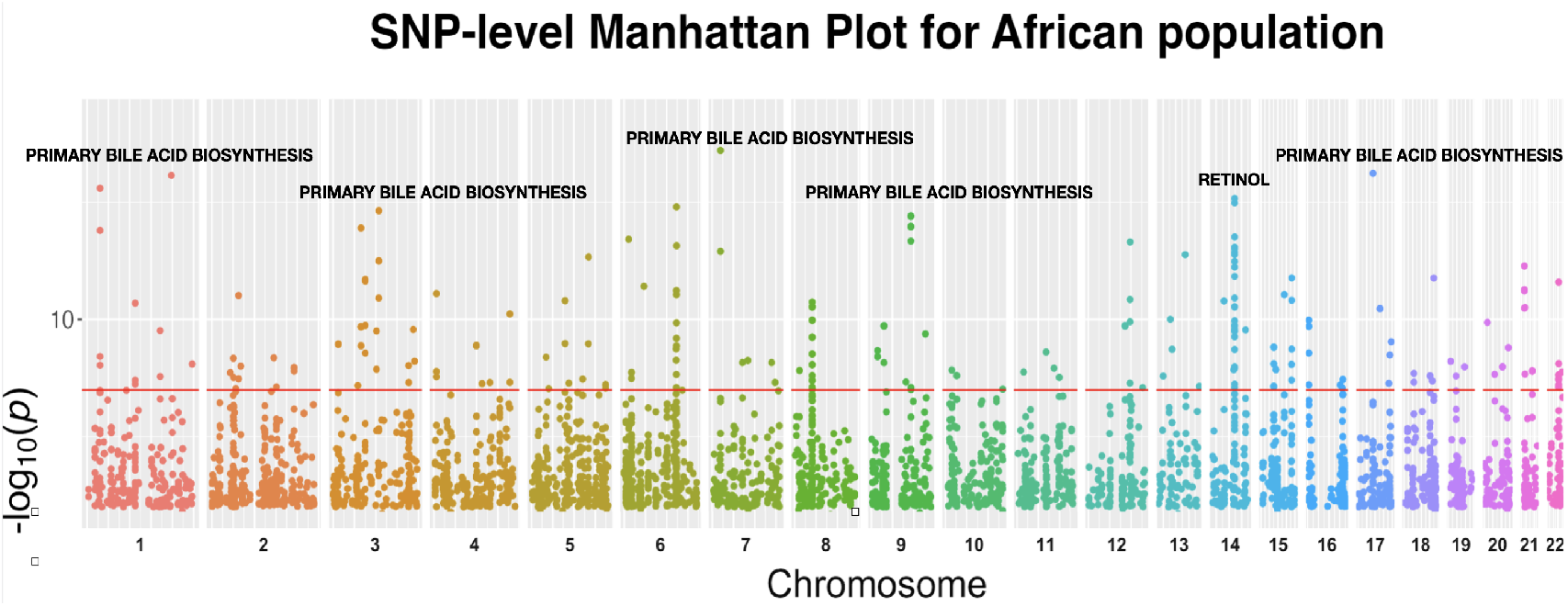
**SNP-level Manhattan plot showing all top SNPs across all traits in the African population. Each dot represents an SNP, with SNPs ordered on the** 𝓍 **-axis according to their genomic position**. *Y* **-axis represents the strength of their association measured as** − log_10_ **transformed** *p***-values starting from** 1 × 10^−6^. **The red line shows the threshold of genome-wide significance** (*p <* 3.2 × 10^−9^).

**Figure 23:**
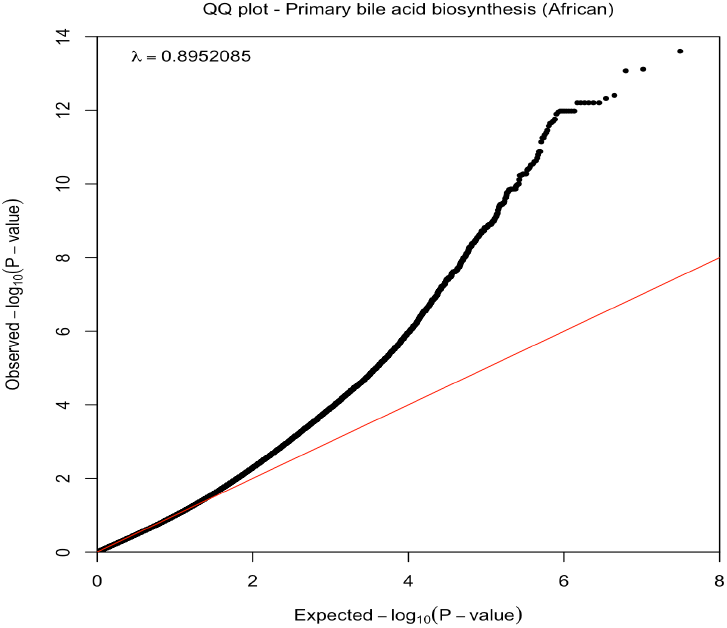
**Quantile–quantile plot showing the distribution of expected** *p***-values under a null model of no significance versus observed** *p***-values for primary bile acid biosynthesis in the African population**.

**Figure 24:**
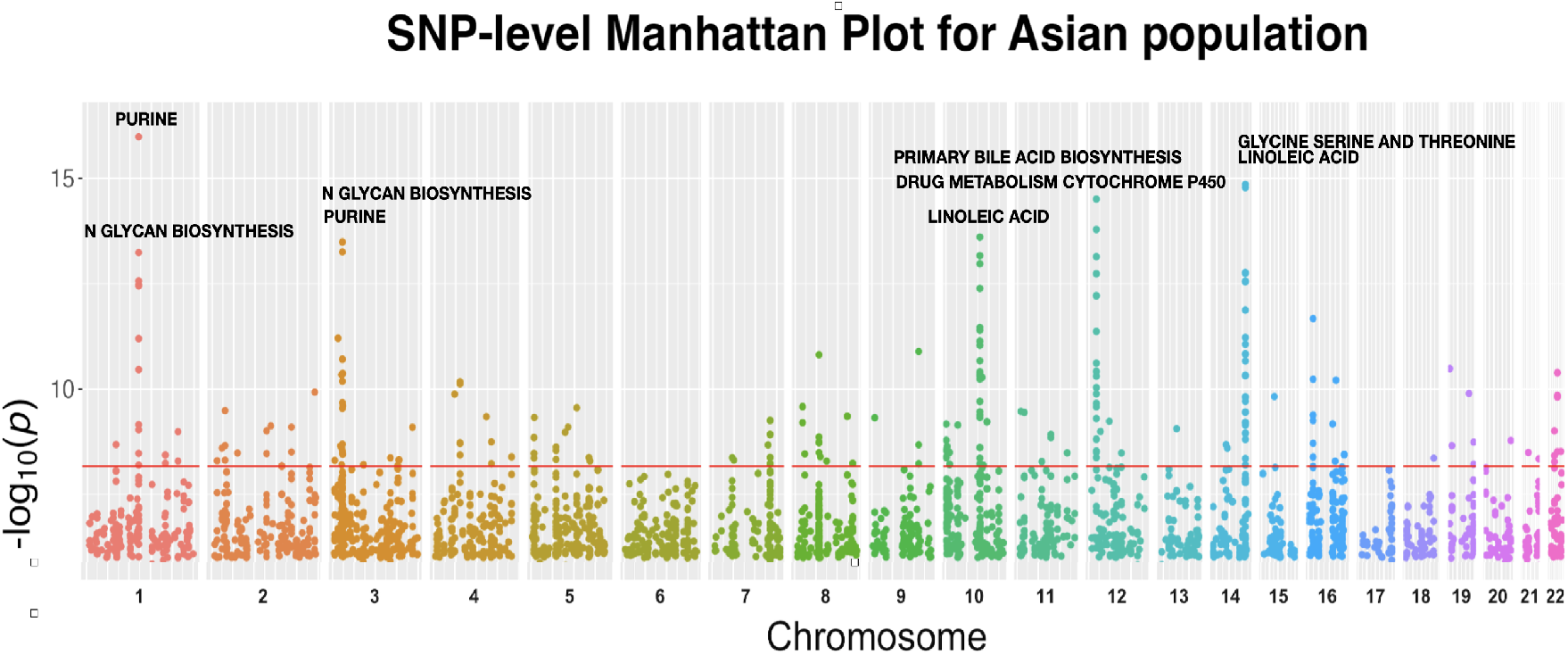
**SNP-level Manhattan plot showing all top SNPs across all traits in the Asian population. Each dot represents an SNP, with SNPs ordered on the** 𝓍 **-axis according to their genomic position**. *Y* **-axis represents the strength of their association measured as** − log_10_ **transformed** *p***-values starting from** 1 × 10^−6^. **The red line shows the threshold of genome-wide significance** (*p <* 6.8 × 10^−9^).

**Figure 25:**
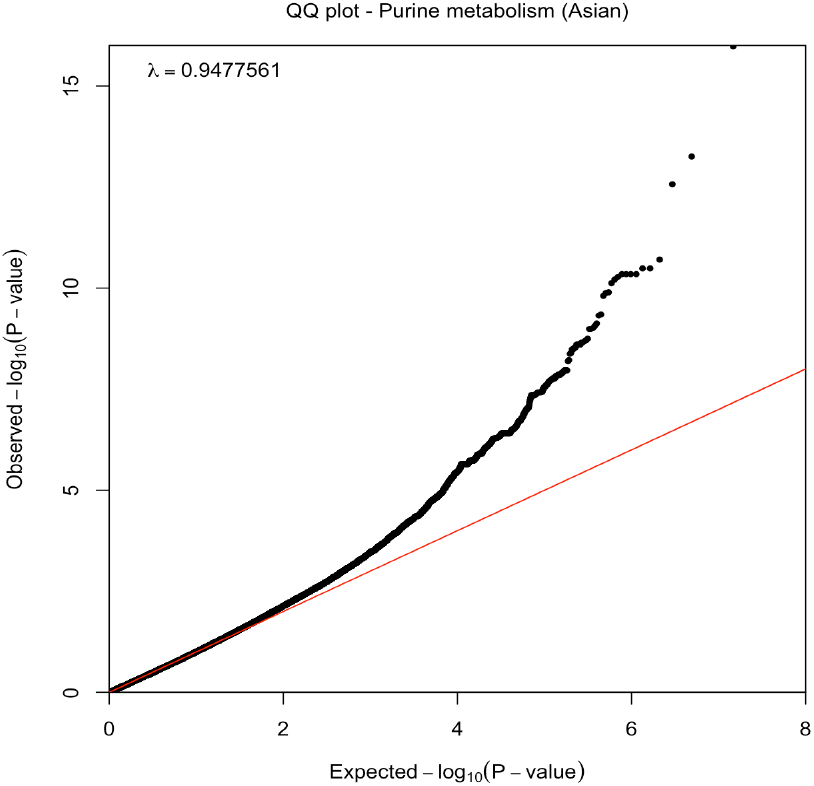
**Quantile–quantile plot showing the distribution of expected** *p***-values under a null model of no significance versus observed** *p***-values for purine metabolism in the Asian population**.

**Figure 26:**
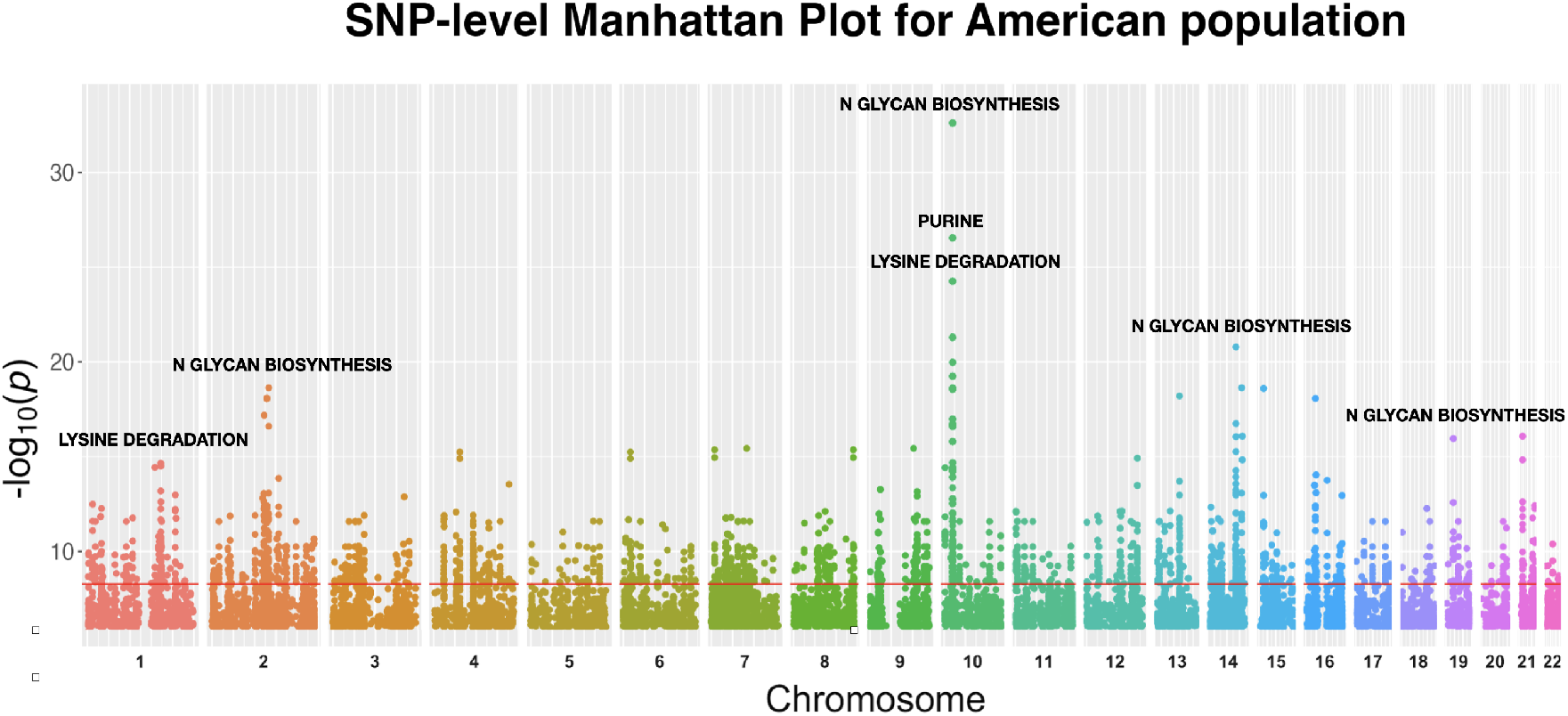
**SNP-level Manhattan plot showing all top SNPs across all traits in the Native American population. Each dot represents an SNP, with SNPs ordered on the** 𝓍 **-axis according to their genomic position**. *Y* **-axis represents the strength of their association measured as** − log_10_ **transformed** *p***-values starting from** 1 × 10^−6^. **The red line shows the threshold of genome-wide significance** (*p <* 5.2 × 10^−9^).

**Figure 27:**
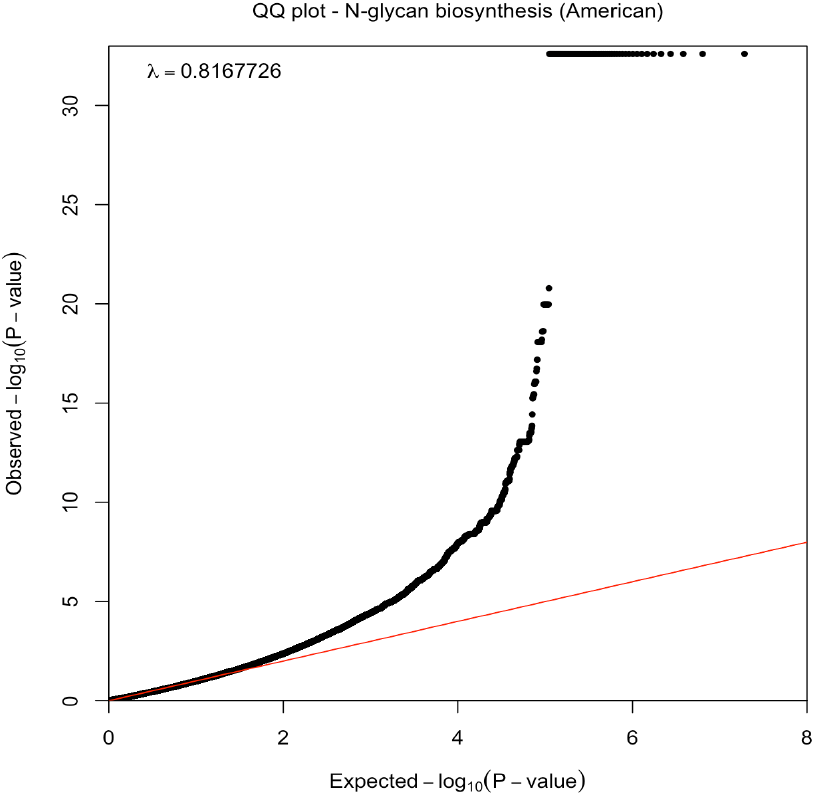
**Quantile–quantile plot showing the distribution of expected** *p***-values under a null model of no significance versus observed** *p***-values for N-glycan biosynthesis in the Native American population**.

**Figure 28:**
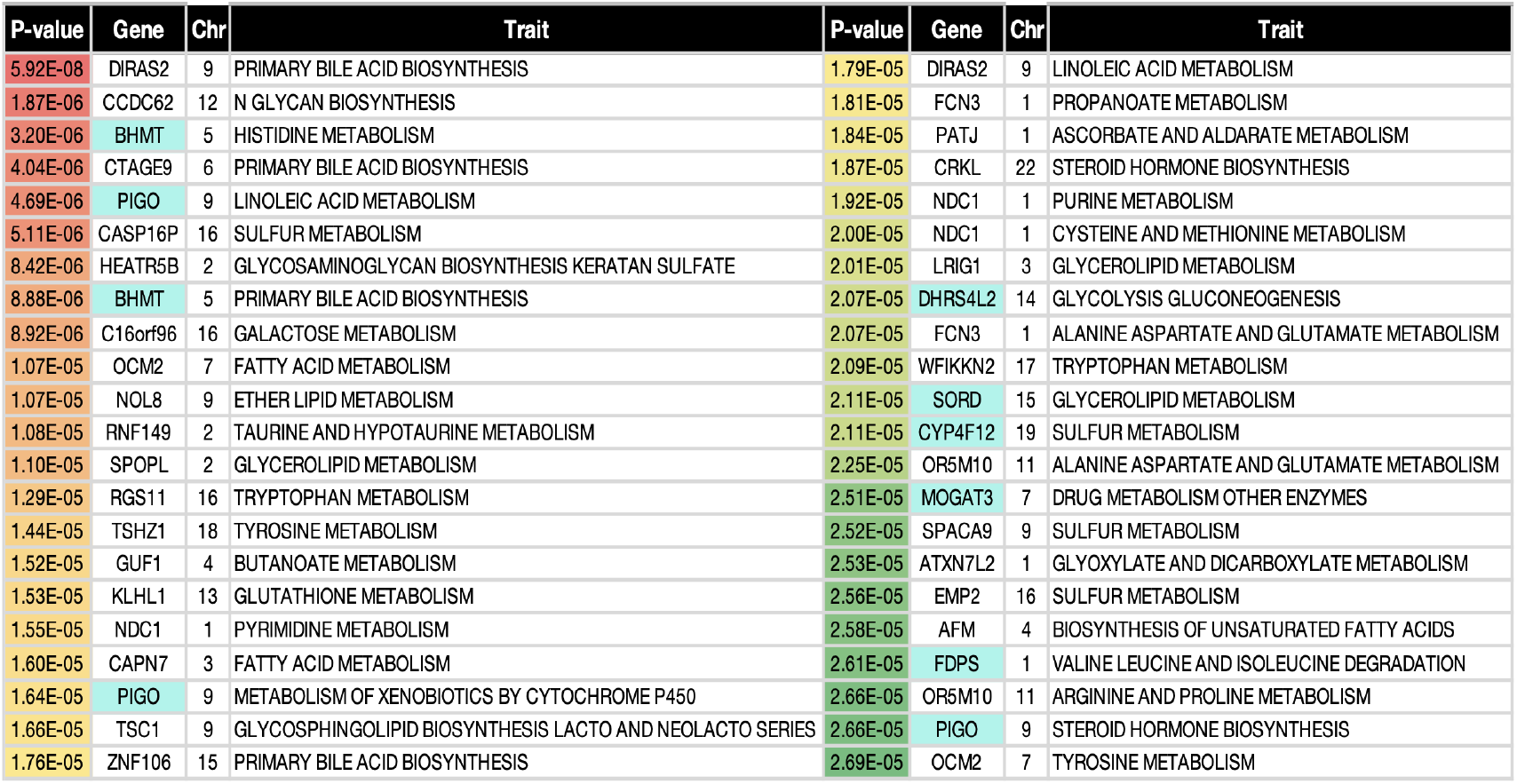
**List of top genes for African ethnic group**. *p***-values tend to be red and green, representing candidate and suggestive genes, respectively. The metabolic genes are highlighted in cyan blue, and the corresponding metabolic traits can be seen in the right-most columns**.

**Figure 29:**
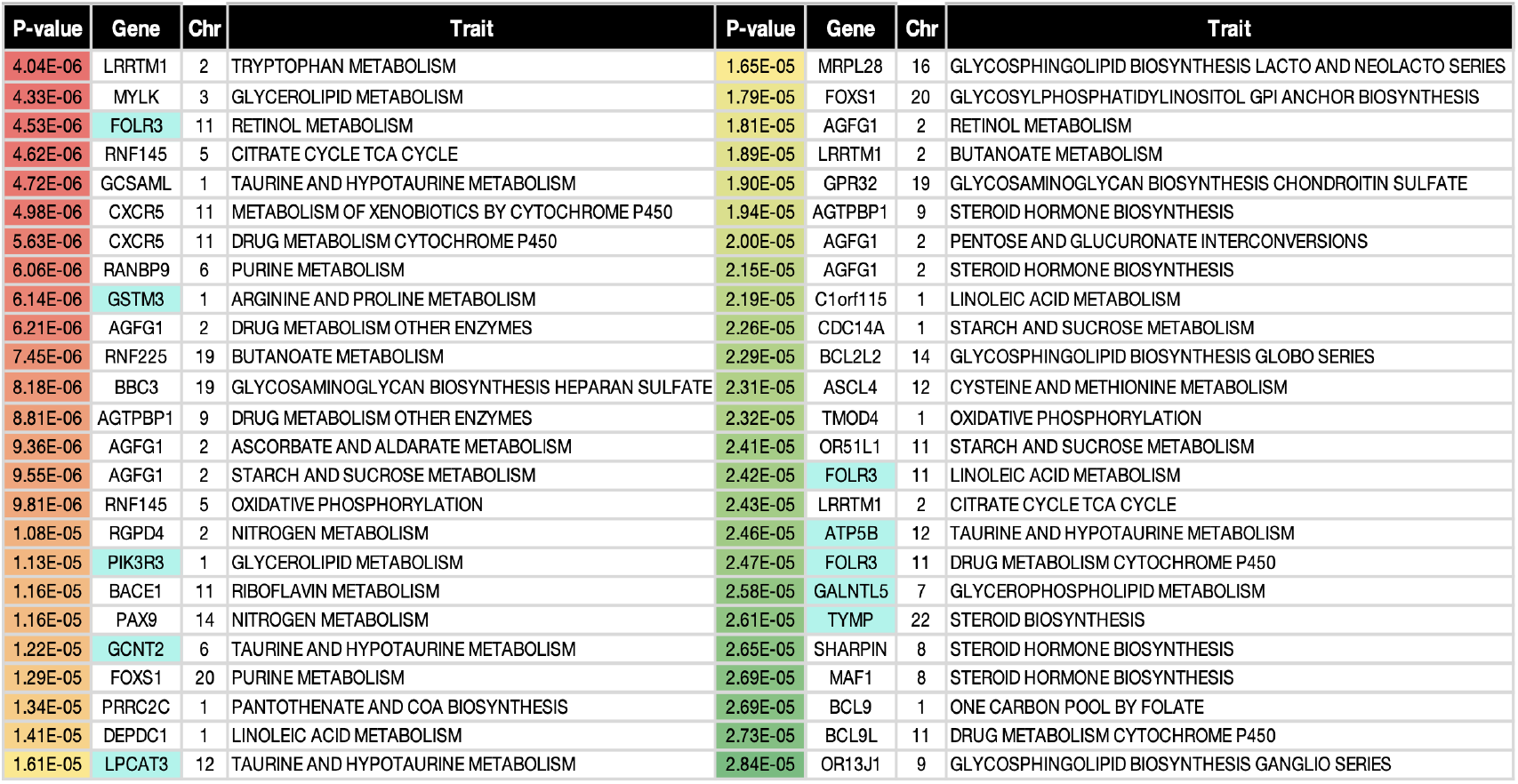
**List of top genes for Asian ethnic group**. *p***-values tend to be red and green, representing candidate and suggestive genes, respectively. The metabolic genes are highlighted in cyan blue, and the corresponding metabolic traits can be seen in the right-most columns**.

**Figure 30:**
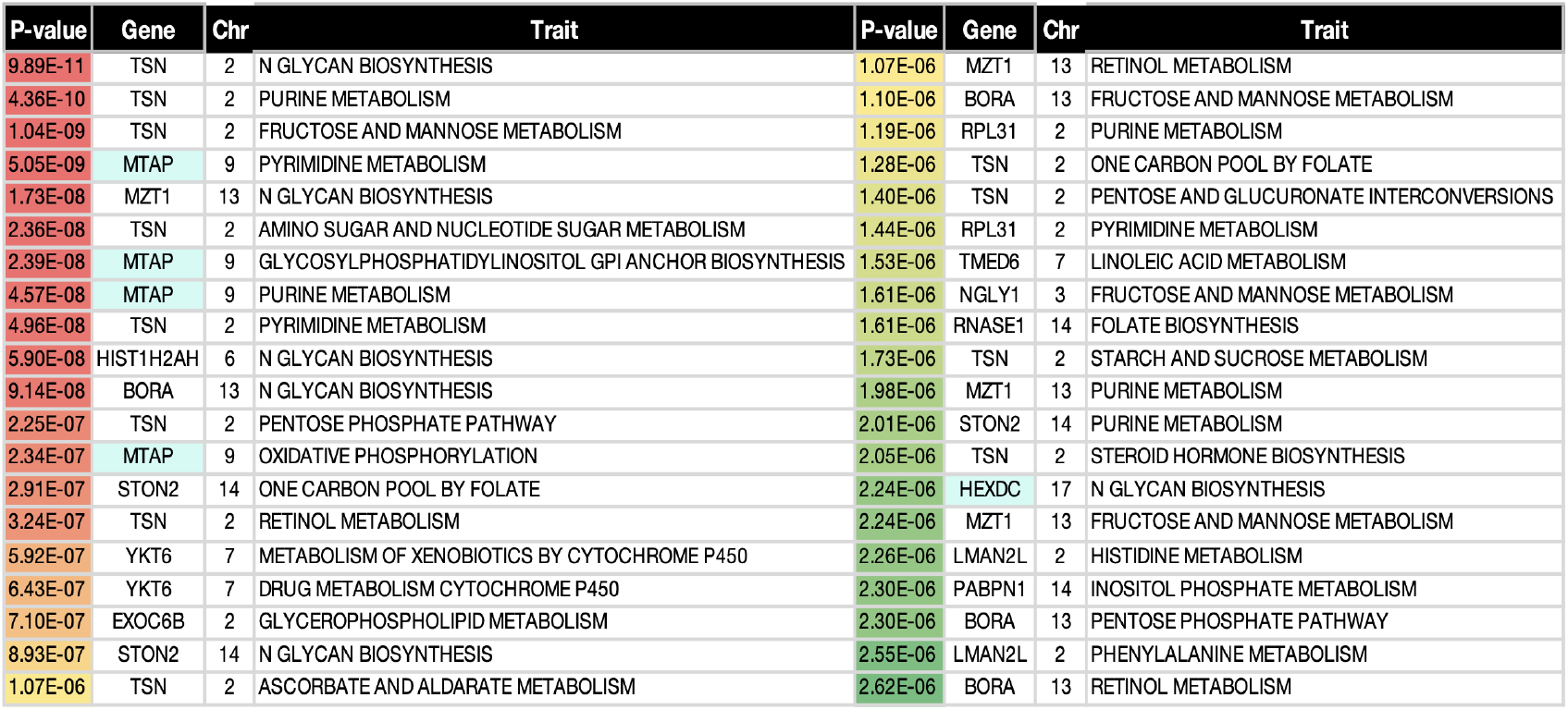
**List of top genes for Native American ethnic group**. *p***-values tend to be red and green, representing candidate genes only for the most to least significant. The metabolic genes are highlighted in cyan blue, and the corresponding metabolic traits can be seen in the right-most columns**.

**Figure 31:**
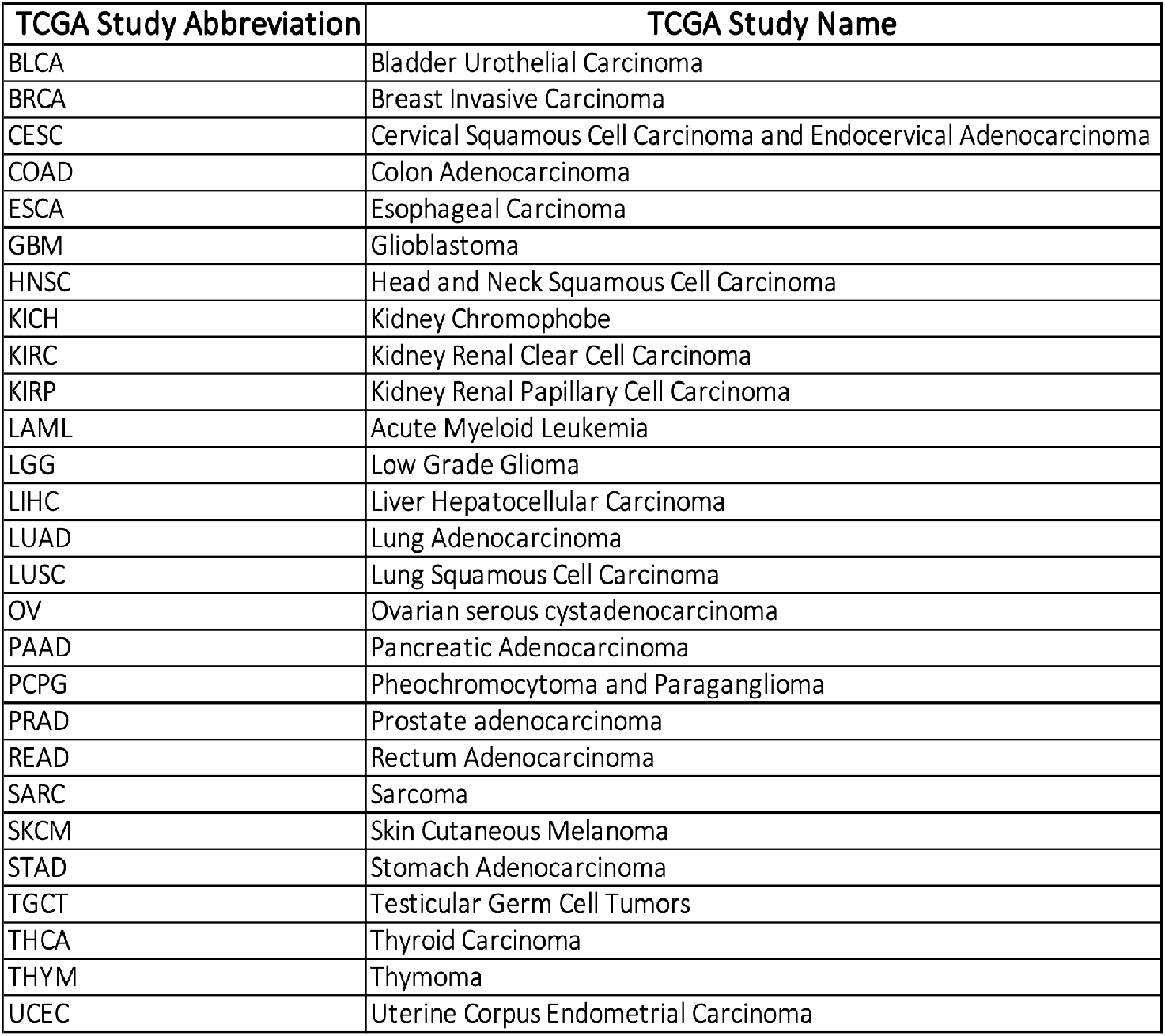
TCGA cancer types included in analysis.

## Footnotes

1 https://gdc.cancer.gov/about-data/publications/pancanatlas

2 https://www.ebi.ac.uk/gwas/

